# ILC3s select for RORγt^+^ Tregs and establish tolerance to intestinal microbiota

**DOI:** 10.1101/2022.04.25.489463

**Authors:** Mengze Lyu, Hiroaki Suzuki, Lan Kang, Fabrina Gaspal, Wenqing Zhou, Jeremy Goc, Lei Zhou, Jordan Zhou, Wen Zhang, JRI Live Cell Bank, Zeli Shen, James G. Fox, Robbyn E. Sockolow, Terri M. Laufer, Yong Fan, Gerard Eberl, David R. Withers, Gregory F. Sonnenberg

## Abstract

Microbial colonization of the mammalian intestine elicits inflammatory or tolerogenic T cell responses, but the mechanisms controlling these distinct outcomes remain poorly understood and accumulating evidence indicates that aberrant immunity to intestinal microbiota is causally associated with infectious, inflammatory, and malignant diseases^1–8^. Here, we define a critical pathway controlling the fate of inflammatory versus tolerogenic T cells that are specific for the microbiota and express the transcription factor RORγt. We profiled all RORγt^+^ immune cells at single cell resolution from the intestine-draining lymph nodes of mice and reveal a dominant presence of Tregs and lymphoid tissue-induced (LTi)-like group 3 innate lymphoid cells (ILC3s), which co-localize at interfollicular regions. These ILC3s have interconverting potential with RORγt^+^ extrathymic Aire-expressing cells, abundantly express major histocompatibility complex class II, and are necessary and sufficient to promote microbiota-specific RORγt^+^ Tregs and prevent their expansion as inflammatory T helper (Th)17 cells. This occurs through ILC3-mediated antigen-presentation, interleukin-2 gradients, and *α*v integrin. Finally, single-cell analyses demonstrate that ILC3 and RORγt^+^ Treg interactions are impaired in inflammatory bowel disease. Our results define a novel paradigm whereby ILC3s positively select for antigen specific RORγt^+^ Tregs, and against Th17 cells, to establish immune tolerance to the microbiota and intestinal health.

## Main

The mammalian gastrointestinal (GI) tract is continuously colonized with microbiota, opportunistic microbes, or pathogens, which induce robust responses that are often dominated by immune cells expressing the lineage-specifying transcription factor RORγt^1–17^. Depending on the adopted fate, these RORγt^+^ cells orchestrate immunity, inflammation or tolerance in the intestine^9–17^ and substantial alterations in RORγt^+^ immune cells occur in multiple chronic human diseases, including inflammatory bowel disease (IBD), HIV infection and cancer^1–17^. Despite these advances, the full spectrum of cellular heterogeneity among RORγt^+^ immune cells, the potential for functional interactions among subsets, and the pathways that are necessary to establish immune tolerance in the context of a complex microbiota remain poorly defined.

### Defining all RORγt^+^ cells in the mLN

RORγt^+^ cell types include T helper (Th)17 cells, regulatory T cells (Tregs), γ*δ* T cells, and group 3 innate lymphoid cells (ILC3s), as well as a few other potential cell types that were only recently characterized^17–19^. To uncover the full spectrum of cellular heterogeneity and examine for pathways impacting tolerance or inflammation to microbes, we performed single-cell RNA-sequencing (scRNA-seq) and profiled all RORγt^+^ cells from the intestine-draining mesenteric lymph nodes (mLN) of healthy RORγt-eGFP reporter mice (**Extended Data Fig. 1a**). T cells are dominant in this tissue, and we therefore sequenced an equal ratio of T cells (GFP^+^TCRβ^+^) to non-T cells (GFP^+^TCRβ^-^) to increase the resolution of rare RORγt^+^ cell types. scRNA-seq revealed 14 distinctive clusters, each of which was annotated based on select marker genes and visualized by uniform manifold approximation and projection (UMAP) (**Fig. 1a**). Two clusters were excluded from future analyses, including cluster 8 that was identified as a doublet population (**Extended Data Fig. 1b**) and cluster 7 that was identified as B cells but could not be verified by RORγt reporter or fate mapping approaches (**Extended Data Fig. 1c, d**), likely representing a contaminate. The remaining clusters all exhibited expression of RORγt and could be divided into T cells versus non-T cells based on expression of *Cd3e* (**Extended Data Fig. 1e, f**). T cell subsets were dominated by RORγt^+^ regulatory T cells (Tregs) defined by expression of *Trac*, *Cd4* and *Foxp3* in cluster 0, 5 and 13, followed by Th17 cells expressing *Trac*, *Cd4* and *Il17a* in cluster 1, and γ*δ* T cells expressing *Trdc* in clusters 4 and 9 (**Fig. 1b, Extended Data Fig. 1g**). Among the non-T cells, nearly all clusters were ILC3s, or ILC3-like cells, based on expression of *Id2* and *Il7r*, including a dominant population of lymphoid tissue-inducer (LTi)-like ILC3s that expressed *Ccr6* and *Cd4* in cluster 2, T-bet^+^ ILC3s that express *Ncr1* in clusters 3 and 12, and a minor population of group 2 innate lymphoid cell (ILC2)-like cells that express high *Gata3* and *Il17rb* in cluster 10 (**Fig. 1c**). Although subsets of conventional type 2 dendritic cells (cDC2s) were reported to express RORγt in the spleen^19^, we could not identify this population in our single cell analyses, and we found minimal expression of RORγt in cDC2s as determined by flow cytometry on reporter or fate mapping mice in the mLN (**Extended Data Fig. 1c, d, h-k**). Instead, the remaining two non-T cell clusters (cluster 6 and 11) were defined by expression of *Aire* (**Fig. 1c**), most closely resembling extra-thymic Aire-expressing cells (eTACs) that several groups defined are RORγt^+^, enriched in lymph nodes during early life developmental windows, and share transcriptional similarities with ILC3s, DCs and thymic epithelial cells^18, 20^.

**Fig. 1.**
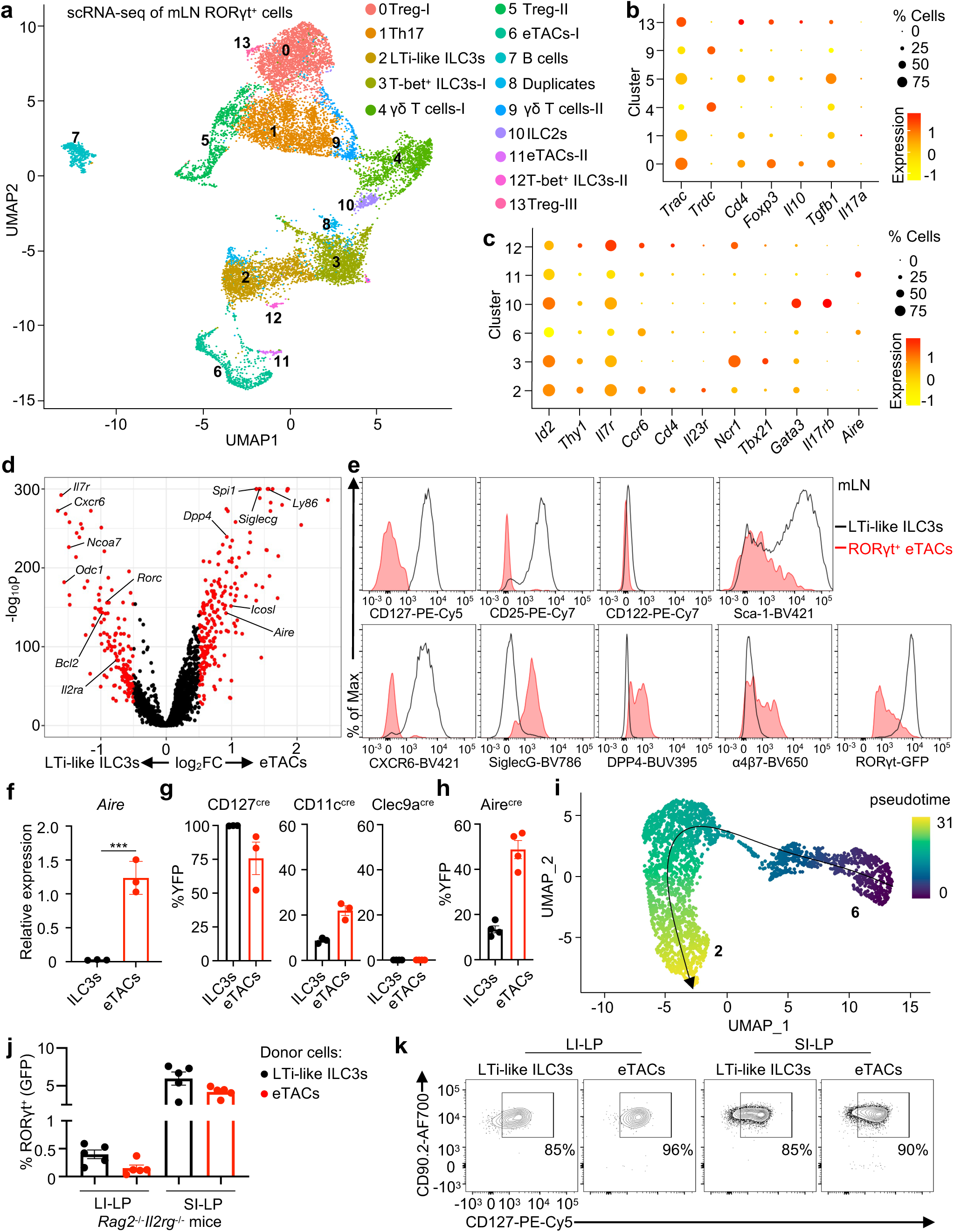
Single-cell resolution of RORγt^+^ adaptive and innate lymphocytes in the mLN. **a**, Uniform manifold approximation and projection (UMAP) of scRNA-seq data from RORγt^+^ cells in mesenteric lymph node (mLN) from healthy RORγt-eGFP reporter mice. Treg: regulatory T cells; LTi-like ILC3: lymphoid tissue inducer-like group 3 innate lymphoid cells; eTACs: extrathymic Aire expressing cells; ILC2: group 2 innate lymphoid cells. TCRβ^+^ GFP^+^ and TCRβ^-^ GFP^+^ cells pooled from 3 mice and 12,948 cells were sequenced. **b**, **c**, Dot plot showing the mean expression (color) of indicated genes in clusters grouped by adaptive lymphocytes (**b**) and innate lymphocytes (**c**), dot size represents the proportion of cells in a cluster with the gene detected. **d**, Volcano plot of differentially expressed genes between cluster 2 and cluster 3 of scRNA-Seq data set. **e**, Histogram examination of indicated genes expression in LTi-like ILC3s (black line) and RORγt^+^ eTACs (red line) from the mLN of RORγt-eGFP reporter mice. **f**, qPCR analysis of *Aire* expression in sort-purified LTi-like ILC3s and RORγt^+^ eTACs from mLN, relative to *Hprt* (n = 3). **g**, **h**, Frequencies of “fate-mapped” LTi-like ILC3 and RORγt^+^ eTACs in the mLN of indicated mice (n = 3-4). **i**, Potential developmental trajectory of indicated clusters inferred by pseudotime analysis plotted on UMAP. **j**, **k**, Graph of the frequencies of RORγt^+^ KLRG1^-^ cells (**j**) with representative flow cytometry plots of CD127^+^CD90^+^ ILC3s (**k**) in large and small intestine of *Rag2^-/-^Il2rg^-/-^* recipient mice (n = 5) with adoptive transferred LTi-like ILC3 and RORγt^+^ eTACs sort purified from RORγt-eGFP reporter mice. LI-LP: large intestine-lamina propria; SI-LP: small intestine-lamina propria. Data in (**f**-**h**, **j**, **k**) are representative of two or three independent experiments, shown as means ± SEM, statistics shown in (**f**) are obtained by unpaired Student’s *t*-test (two-tailed), ***p<0.001.

### RORγt^+^ eTACs can interconvert with ILC3s

Given this heterogeneity in innate RORγt^+^ immune cells, we next sought to understand the phenotype and relationships of ILC3s versus RORγt^+^ eTACs. Differential analysis of LTi-like ILC3s (cluster 2) and the dominant RORγt^+^ eTACs (cluster 6) revealed distinct gene signatures (**Fig. 1d**). Further validation revealed that LTi-like ILC3s are defined by high expression of CD127, CD25, CD122, Sca-1 and CXCR6, while RORγt^+^ eTACs are defined by high expression of SiglecG, DPP4 and integrin *α*4*β*7 (**Fig. 1e, Extended Data Fig. 2a-c**). Notably, LTi-like ILC3s express significantly higher levels of RORγt than eTACs (**Fig. 1d, e, Extended Data Fig. 2c**). We further developed a novel gating strategy to isolate these cell types and confirmed that expression of *Aire* is exclusive to RORγt^+^ eTACs and not observed in ILC3s (**Fig. 1f, Extended Data Fig. 3a**). Notably, RORγt^+^ eTACs did not exhibit expression of previously defined Aire-dependent tissue-specific antigens, but rather expressed several genes associated with a transient-amplifying Aire^+^ thymic epithelial cell^21, 22^ (**Supplementary Table 1**). A previous study inferred that ILC3s have the potential to convert into RORγt^+^ eTACs *in vitro* following RANK-RANKL stimulation^18^, but our identified phenotype of RORγt^+^ eTACs more closely resembled recently described early ILC progenitors (EILP) that are CD127^-^, CD90.2^-^, CD25^-^, and *α*4*β*7^+^ cells^23, 24^. This provokes fundamental questions about the *in vivo* interconverting potential of these two cell types. We determined that a majority of RORγt^+^ eTACs fate-mapped positive for IL-7R-cre, despite lacking CD127 staining, and exhibited limited fate-mapping for CD11c-cre or Clec9a-cre, which is comparable to LTi-like ILC3s (**Fig. 1g, Extended Data Fig. 3a, b**). In addition, RORγt^+^ eTACs fate-mapped positive for Aire-cre, and so do a minor proportion of LTi-like ILC3s in the mLN (**Fig. 1h, Extended Data Fig. 3b**). Lineage trajectory analyses of our single cell data set suggest that RORγt^+^ eTACs have the potential to give rise to LTi-like ILC3s (**Fig. 1i, Extended Data Fig. 3c**). To test this model, we performed an adoptive cell transfer of sort purified LTi-like ILC3s or RORγt^+^ eTACs into recipient *Rag2^-/-^Il2rg^-/-^* mice and determined that both had the potential to expand and reconstitute as conventional CD90.2^+^CD127^+^ ILC3s in the intestines (**Fig. 1j, k, Extended Data Fig. 4a, b**). In these contexts, RORγt^+^ eTAC-derived ILC3s exhibit staining for CCR6 and robustly fate-map for a history of Aire-expression but did not interconvert into NK cells, ILC1s or ILC2s (**Extended Data Fig. 4c-f**). Collectively, these data define the full spectrum of innate and adaptive immune cells that express the lineage-specifying transcription factor RORγt in the mLN of healthy mice, revealing a dominance of RORγt^+^ Tregs and LTi-like ILC3s, as well as determining that RORγt^+^ eTACs have the potential to interconvert into LTi-like ILC3s in certain contexts.

### RORγt^+^ Tregs require MHCII^+^ ILC3s

We next sought to define functional interactions between innate and adaptive RORγt^+^ cells in mLN by examining for their spatial proximity in the interfollicular zone, an area we previously found to contain a majority of ILC3s and other RORγt^+^ T cells^25, 26^. Indeed, ILC3s were rare in B cell follicles and T cell zones, but robustly present in interfollicular zones, which contrasts to more diffuse localization patterns of CD11c^+^ cells (**Extended Data Fig. 5**). Strikingly we observed that CD3^-^RORγt^+^CD127^+^ ILC3s were adjacent to CD3^+^RORγt^+^FoxP3^+^ Tregs in the interfollicular zones, where a substantial portion of total RORγt^+^ Tregs were associated with ILC3s (**Fig. 2a, Extended Data Fig. 5**). CD103^+^ dendritic cells (DCs) were proposed to induce the differentiation of RORγt^+^ Treg from naïve T cells^27–30^, however both CD103^+^ and CD103^-^ DCs were recently reported to redundantly promote peripheral regulatory T cells differentiation^31^, indicating other cell types regulate this process. Our previous work demonstrated LTi-like ILC3s limit microbiota-specific Th17 cells via major histocompatibility complex class II (MHCII) and a process termed “intestinal selection”^26, 32^. Therefore, we next interrogated whether LTi-like ILC3s impact RORγt^+^ Treg through MHCII and observed that LTi-like ILC3s abundantly express MHCII protein and transcripts in the mLN of healthy mice (**Fig. 2b, Extended Data Fig. 6a**). Further, MHCII^ΔILC3^ mice that were generated by crossing *H2-Ab1^fl/fl^* mice with *Rorc^cre^* mice^32^ exhibited a highly selective deletion of MHCII in ILC3s relative to other immune cells in the draining mLN and large intestine (**Fig. 2c**). In mice lacking ILC3-specific MHCII, we identified a striking and significant reduction in the frequency of RORγt^+^ Tregs in both mLN and large intestine relative to littermate control mice (**Fig. 2d-g**). In addition, as we previously reported, deletion of ILC3-specific MHCII results in significantly increased total CD4^+^ T cells and Th17 cells within the large intestine, with a significant decrease in numbers of RORγt^+^ Tregs (**Extended Data Fig. 6b-e**). These results were specific to MHCII^+^ LTi-like ILC3s, as deletion of MHCII by targeting T cells with CD4-cre or ILC2s with IL-5-cre, and deletion of RORγt by targeting T-bet^+^ ILC3s with Ncr1-cre, did not impact the frequency of RORγt^+^ Tregs or other CD4^+^ T cell subsets in the large intestine (**Extended Data Fig. 6f-k**).

**Fig. 2.**
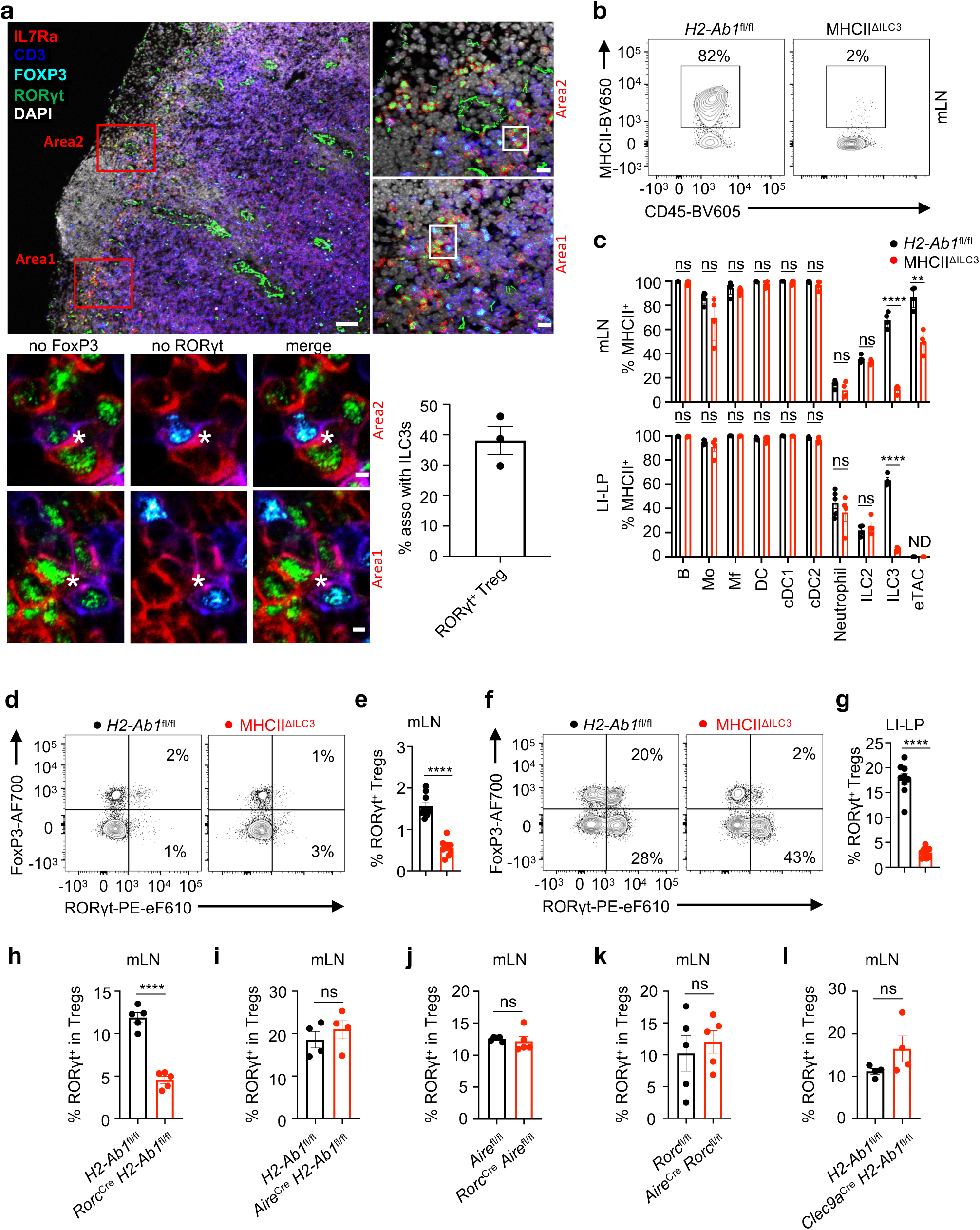
LTi-like ILC3s select for microbiota specific RORγt^+^ Tregs via MHCII. **a**, Tile-scanned (top left) and magnified (top right) images of mLN stained for expression of IL7Rα (red), CD3 (blue), FOXP3 (cyan), RORγt (green) and DAPI (grey). Serial sections (bottom left) from mLN as shown in (top left) showing the same interfollicular area stained for ILC3s (CD3^-^ IL7Rα^+^RORγt^+^) and RORγt^+^ Treg (CD3^+^IL7Rα^-^FOXP3^+^RORγt^+^), white asterisk indicates the association between ILC3s and RORγt^+^ Treg. Left panels are without FoxP3 staining, middle panels are without RORγt staining, and right panels are a merge. Scale bar: 50 μm (top left), 10 μm (top right) and 2 μm (bottom left). Quantification (bottom right) of percent of RORγt^+^ Treg in association with ILC3 in the interfollicular zone of the mLN (n = 3). **b**, Representative flow cytometry plots of the frequency of MHCII on LTi-like ILC3s in mLN of *H2-Ab1*^fl/fl^ and MHCII^ΔILC3^ mice. **c**, Quantification of MHCII expression on major MHCII-expressing cells from mLN and LI-LP of *H2-Ab1*^fl/fl^ and MHCII^ΔILC3^ mice (n = 4-5). Mo: monocytes; Mf: macrophages; DC: dendritic cells; cDC: conventional dendritic cells; ILC: innate lymphoid cells; eTACs: extrathymic Aire expressing cells. ND, not detected. **d-g**, Representative flow cytometry plot of the frequency (% of CD4^+^ T cells) (**d**, **f**) and quantification (**e**, **g**) of RORγt^+^ Tregs (CD45^+^CD3^+^CD4^+^FoxP3^+^RORγt^+^) in mLN and LI-LP of *H2-Ab1*^fl/fl^ and MHCII^ΔILC3^ mice (n = 9). **h**-**l**, Quantification of RORγt^+^ Tregs (% of Tregs) in mLN of *H2-Ab1*^fl/fl^ and MHCII^ΔILC3^ mice (n = 5) (**h**), *H2-Ab1*^fl/fl^ and *Aire*^cre^ x *H2-Ab1*^fl/fl^ mice (n = 4) (**i**), *Aire*^fl/fl^ and *Rorc*^cre^ x *Aire*^fl/fl^ mice (n = 5) (**j**), *Rorc*^fl/fl^ and *Aire*^cre^ x *Rorc*^fl/fl^ mice (n = 5) (**k**) and *H2-Ab1*^fl/fl^ and *Clec9a*^cre^ x *H2-Ab1*^fl/fl^ mice (n = 4) (**l**). Data in **d**-**g** are pooled from two independent experiments. Data in **h**-**l** are representative of two independent experiments. Data are shown as mean ± SEM, statistics shown in (**c**) are obtained by multiple unpaired *t*-test, statistics shown in (**e**, **g**, **h**-**l**) are obtained by unpaired Student’s *t*-test (two-tailed), ns, not significant; **p<0.01, ****p<0.0001.

RORγt^+^ eTACs also express MHCII as previously shown^21^, but *Rorc^cre^* only modestly targeted deletion of MHCII on RORγt^+^ eTACs and this is likely due to their significantly lower expression of RORγt relative to ILC3s (**Fig. 1d, e, Fig. 2c, Extended Data Fig. 6a**). Further, more specific targeting of RORγt^+^ eTACs also indicated a redundant role for this population in modulating RORγt^+^ Tregs as comparable frequencies were present in *Rorc^cre^* x *Aire^fl/fl^* mice, *Aire^cre^* x *Rorc^fl/fl^* mice, and *Aire^cre^* x *H2-Ab1^fl/fl^* mice relative to littermate controls (**Fig. 2h-k, Extended Data Fig. 6l-p**). LTi-like ILC3s were also comparable in *Aire^cre^* x *Rorc^fl/fl^* relative to littermates (**Extended Data Fig. 6n**), indicating that RORγt^+^ eTACs are redundant to maintain this population in homeostasis. RORγt^+^ Tregs were also present at comparable frequencies in *Clec9a^cre^* x *H2-Ab1^fl/fl^* mice relative to literate controls, indicating a redundancy of cDCs (**Fig. 2l, Extended Data Fig. 6q**). Finally, an *in vitro* co-culture system revealed that RORγt^+^ Tregs significantly increased in frequency and cell number when in the presence of LTi-like ILC3s, as well as exhibited reduced Bim, increased Nur77 staining and minor changes in cell death at this time point (**Extended Data Fig. 7a-e**), indicating that LTi-like ILC3s are sufficient to support RORγt^+^ Tregs.

### ILC3s select for microbiota specific Tregs

RORγt^+^ T cells develop in response to antigens derived from microbiota that colonize the mammalian intestine. For example, studies demonstrated that segmented filamentous bacteria (SFB) promote antigen-specific Th17 cells^33^, while *Helicobacter hepaticus* (*H. hepaticus*) promote antigen-specific RORγt^+^ Tregs in the intestine of wild-type mice^34^. Therefore, we next examined whether MHCII^+^ ILC3s regulate both Th17 cells and RORγt^+^ Tregs that recognize distinct microbiota-derived antigens in the intestine. To accomplish this, we transferred congenically-marked naïve T cells from SFB-specific TCR transgenic (7B8) mice and *H. hepaticus*-specific TCR transgenic mice (HH7-2) into recipients that were colonized with both microbes (**Extended Data Fig. 8a**). Two weeks post-transfer, we analyzed the donor T cell populations and observed comparable upregulation of CD44 in recipient littermate controls and MHCII^ΔILC3^ mice (**Extended Data Fig. 8b, c**). This indicates that ILC3s are not required to prime antigen-specific T cells and is consistent with the absence of ILC3s from T cell zones of lymph nodes (**Extended Data Fig. 5**)^26^. SFB-specific T cells differentiated into Th17 cells in littermate control mice and a significantly expanded Th17 cell population was observed in mice lacking ILC3-specific MHCII (**Fig. 3a, b**). The impairment of RORγt^+^ Tregs is insufficient to drive expansion of effector Th17 cells, such as the SFB-specific populations, as comparable intestinal Th17 cells were observed in *Foxp3^cre^* x *Rorc^fl/fl^* mice (**Extended Data Fig. 8d**). In contrast, we observed that intestinal *H. hepaticus*-specific T cells failed to express FoxP3 in mice lacking ILC3-specific MHCII, and rather adopted a RORγt^+^ Th17 cell phenotype relative to those transferred to littermate controls (**Fig. 3c, d**). In the absence of ILC3-specific MHCII, the *H. hepaticus*-specific T cell also upregulated T-bet and IFN-γ, as well as co-produced significantly more IFN-γ and IL-17A (**Fig. 3e-h**). We also observed a significant impairment in the differentiation of *H. hepaticus*-specific RORγt^+^ Tregs when ILC3s were targeted in *Il22^cre^* x *H2-Ab1^fl/fl^* mice, but in this context, there was only a partial but selective deletion of MHCII on ILC3s in the mLN (**Extended Data Fig. 8e-g**). These data demonstrate that ILC3-specific MHCII is necessary to limit the expansion of microbiota specific effector Th17 cells as previously described^26, 32^, while also simultaneously enforcing microbiota specific RORγt^+^ Treg populations and preventing their ability to expand as pro-inflammatory Th17 cells.

**Fig. 3.**
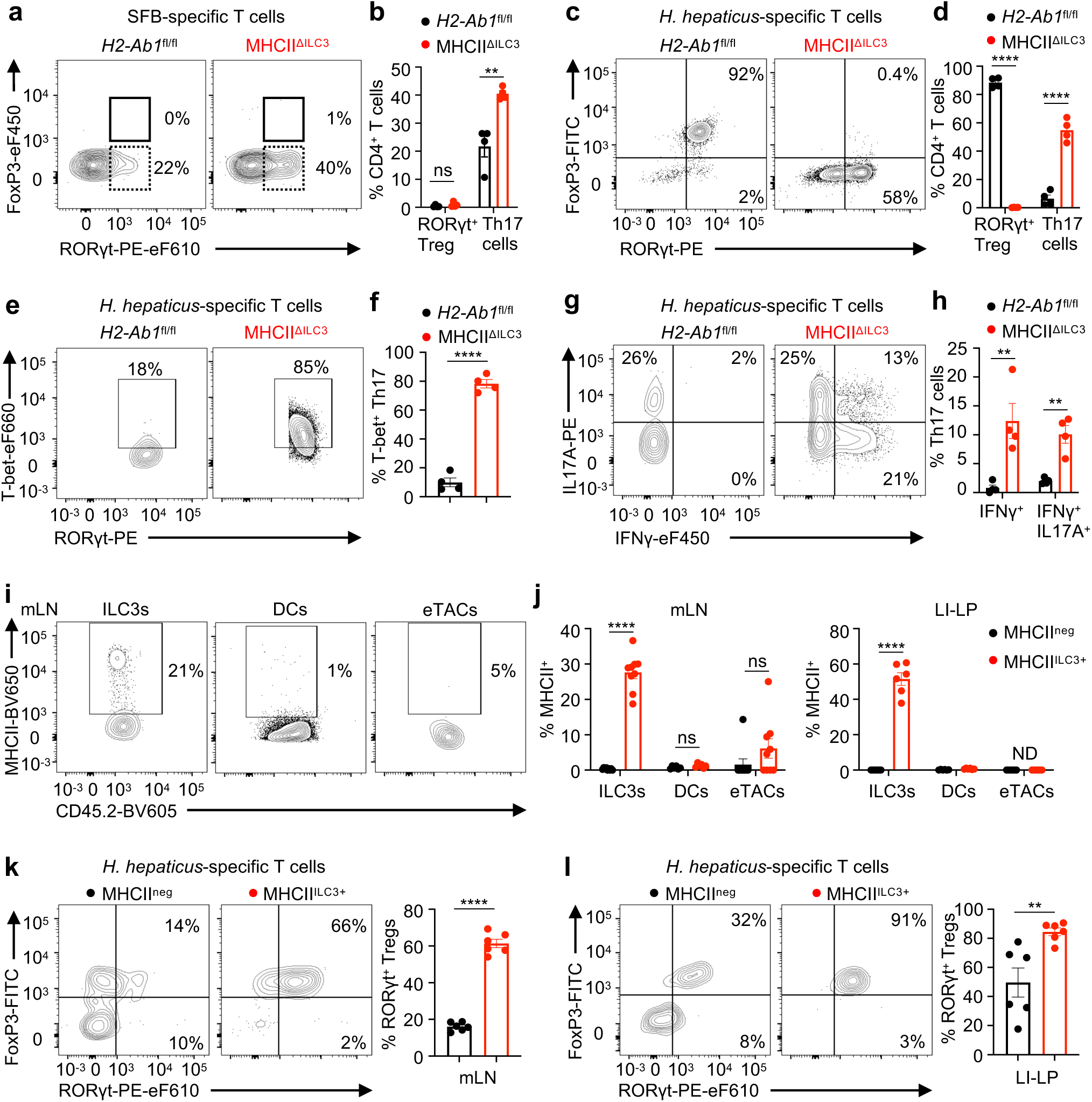
MHCII+ ILC3s are sufficient for selection of microbiota specific RORγt^+^ Tregs. **a**, Frequency of RORγt^+^ Tregs (RORγt^+^FoxP3^+^ among SFB or Hh-specific CD4^+^ T cells) and Th17 cells (RORγt^+^FoxP3^-^ among SFB or Hh-specific CD4^+^ T cells) were analyzed in Peyer’s patch (PP) for SFB7B8 (**a**, **b**, CD45.1^-^CD90.1^+^CD4^+^ T cells) and in colon for HH7-2 (**c**, **d**, CD45.1^+^CD90.1^-^CD4^+^ T cells) of *H2-Ab1*^fl/fl^ and MHCII^ΔILC3^ mice (n = 4); (**a**, **c**) flow cytometry plot; (**b**, **d**) quantification of frequency. **e**, **f**, Representative flow cytometry plots of the frequency (T-bet^+^ among Hh-specific RORγt^+^FoxP3^-^ Th17 cells) (**e**) and quantification (**f**) of T-bet^+^ Hh-specific Th17 in LI-LP of *H2-Ab1*^fl/fl^ and MHCII^ΔILC3^ mice as shown in (**c**) (n = 4). **g**, **h**, Representative flow cytometry plots of the frequency (IL17A^+^ and/or IFNγ^+^ among Hh-specific RORγt^+^FoxP3^-^ Th17 cells) (**g**) and quantification (**h**) of IFNγ^+^ and IFNγ^+^IL17A^+^ Hh-specific Th17 in LI-LP of *H2-Ab1*^fl/fl^ and MHCII^ΔILC3^ mice as shown in (**c**) (n = 4). **i**, **j**, Representative flow cytometry plots of the frequency (**i**) and quantification (**j**) of MHCII expression on ILC3s, DCs and eTACs (gated as CD127^-^CD90^-^) in mLN and LI-LP of MHCII^neg^ and MHCII^ILC3+^ mice (n = 6-9). ND, not detected. **k**, **l**, Representative flow cytometry plots of the frequency and quantification of RORγt^+^ Tregs (RORγt^+^FoxP3^+^ among Hh-specific CD4^+^ T cells) were analyzed for HH7-2 T cells in mLN (**k**) and LI-LP (**l**) of MHCII^neg^ and MHCII^ILC3+^ mice (n = 6). Data in **a**-**h** are representative of two independent experiments. Data in **i**-**l** are pooled from two or three independent experiments. representative of two independent experiments Data are shown as mean ± SEM, statistics are obtained by unpaired Student’s *t*-test (two-tailed), ns, not significant; **p<0.01, ****p<0.0001.

We also interrogated whether MHCII^+^ ILC3s are sufficient to promote microbiota specific RORγt^+^ Tregs by utilizing mice in which MHCII expression is restricted to only ILC3s^27^. Revisiting *H2-Ab1-*STOP*^fl/fl^* mice (MHCII^neg^) and those crossed to *Rorc^cre^* mice (MHCII^ILC3+^) revealed a robust expression of MHCII restricted only to ILC3s, but not on cDCs in the mLN and large intestine, as well as a lack of MHCII on RORγt^+^ eTACs in the mLN that is likely due to the limited expression of *Rorc* (**Fig. 3i, j, Extended Data Fig. 8h**). We next transferred congenically-marked naïve T cells from *H. hepaticus*-specific TCR transgenic mice into recipients that were colonized with *H. hepaticus* as above. Two weeks post-transfer, we analyzed the donor T cell populations and observed comparable upregulation of CD44 and downregulation of CD62L in recipient MHCII^neg^ and MHCII^ILC3+^ mice (**Extended Data Fig. 8i**). This suggest that even with the lack of MHCII in this MHCII^neg^ mouse model, there remains some endogenous priming of naïve T cells. However, when analyzing the fate of these T cells, it was clear that MHCII expression on ILC3s is sufficient for *H. hepaticus*-specific T cells to efficiently adopt a RORγt^+^ Treg fate in both the mLN and large intestine (**Fig. 3k, l**). These results collectively demonstrate that MHCII^+^ ILC3s are necessary and sufficient to critically select for the fates of RORγt^+^ T cells by simultaneously promoting Tregs and limiting Th17 cells of distinct antigen-specificities.

### ILC3s support RORγt^+^ Tregs via Itgav

We next mechanistically examined how MHCII^+^ ILC3s promote RORγt^+^ Tregs. Analyses of the remaining RORγt^+^ Treg population in MHCII^ΔILC3^ mice revealed a significant increase in Bim but comparable levels of Ki-67 (**Extended Data Fig. 8j, k**), suggesting there are alterations in cell survival but not proliferation. Defined receptor-ligand analyses in our scRNA-seq data demonstrated that MHCII^+^ LTi-like ILC3s may interact with RORγt^+^ Tregs through cytokine-cytokine receptors or the integrin Itgav-mediated processing of transforming growth factor (TGF)-β (**Fig. 4a**). ILC3-derived IL-2 was not involved in this process, as ablation of IL-2 in all RORγt^+^ cells did not impact the presence of RORγt^+^ Tregs (**Extended Data Fig. 8l**). We also previously defined that ILC3-mediated sequestration of IL-2 mechanistically contributed to their ability to limit effector T cell responses^27^, but analyses of IL-2 binding and CD25 levels revealed that RORγt^+^ Tregs were significantly more efficient at competing for this survival cytokine over naïve or effector T cell populations (**Extended Data Fig. 8m**). Therefore, we next explored integrins that mediate processing of TGF-β, as prior reports identified a critical role of DC-mediated processing of TGF-β by integrin αvβ8 in the support of Tregs^35–37^ and that loss of TGFβR on Tregs results in upregulation of Bim^38^. We explored what β-integrins pair with Itgav on ILC3s and found moderate levels of *Itgb1*, *Itgb3*, and *Itgb5* expression on ILC3s in the mLN and large intestine, but a lack of *Itgb6* and *Itgb8* (**Fig. 4a, b**). Utilizing our *in vitro* co-culture system, we found that blockade of Itgav or Itgb3 significantly abrogated ILC3-mediated support of RORγt^+^ Tregs and suppression of Th17 cells (**Fig. 4c**). This indicates that ILC3s interact with RORγt^+^ Tregs in part through αvβ3 integrin, which has previously been linked to processing and activation of TGF-β but also several other extracellular ligands^39^. Consistent with this, we found that ILC3s in the mLN and intestine stained highly for integrin αv (Itgav), and that this could be successfully deleted by crossing *Itgav^fl/fl^* mice with *Rorc^cre^* mice, while Itgav remained intact on RORγt^+^ eTACs in these mice likely due to the limited expression of *Rorc* (**Fig. 4d, Extended Data Fig. 8n**). Consistent with the role we identified for MHCII^+^ ILC3s, we observed a significant reduction in the frequency of RORγt^+^ Tregs and a significant increase in the frequency of Th17 cells in both the mLN and large intestine of *Rorc^cre^ Itgav^fl/fl^* mice relative to littermate control mice (**Fig. 4e**). In addition, *H. hepaticus*-specific T cells failed to robustly differentiate into RORγt^+^ Tregs, and rather expanded as Th17 cells in both the mLN and large intestine of *Rorc^cre^ Itgav^fl/fl^* mice relative to littermate controls (**Fig. 4f, Extended Data Fig. 8o**). Taken together, these data indicate that MHCII^+^ ILC3s select for antigen specific RORγt^+^ Tregs, and that this occurs in part by ILC3 expression of the αvβ3 integrin, which can promote the processing of TGF-β or interact with other extracellular ligands that could be important in this process.

**Fig. 4.**
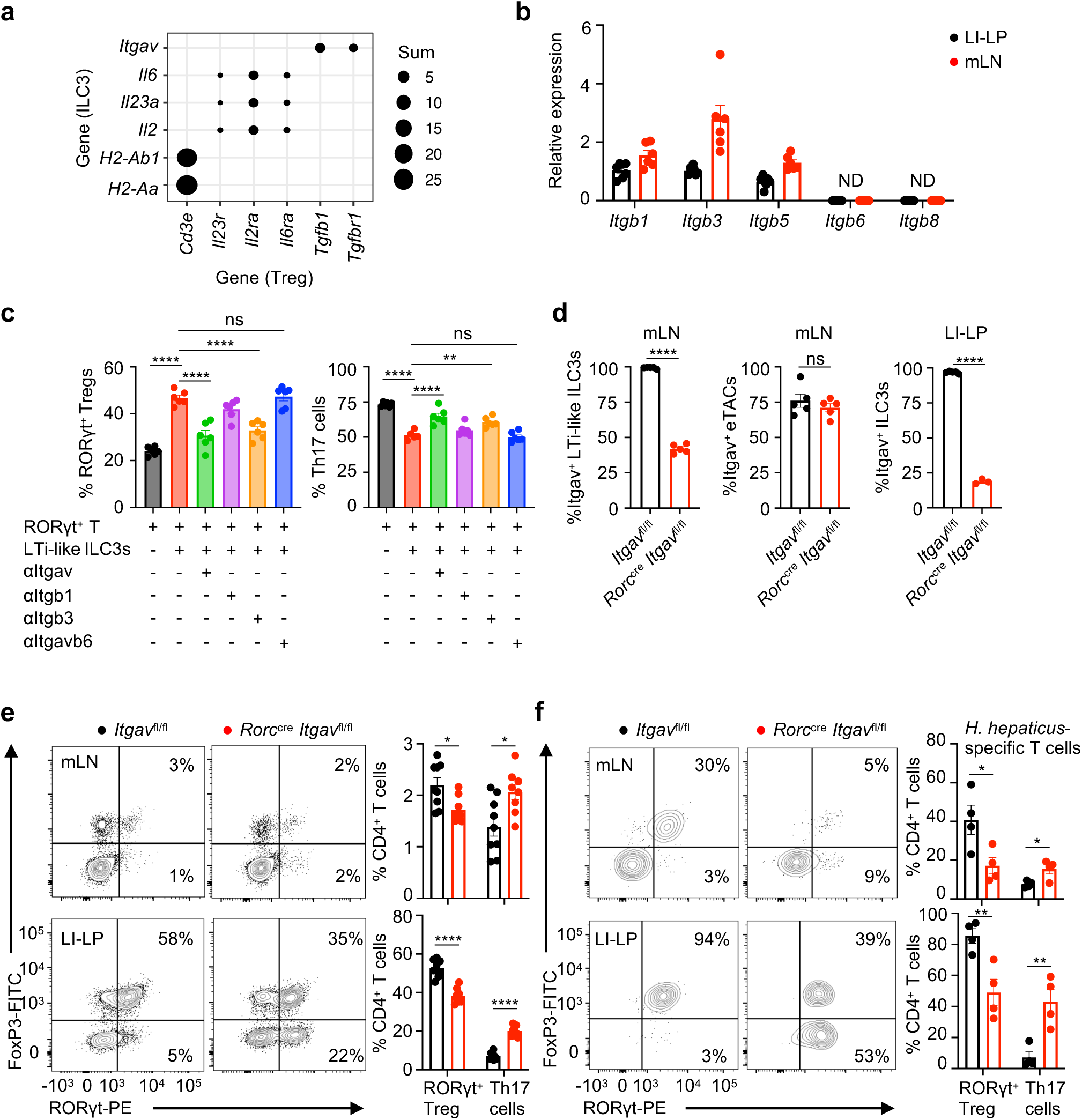
Itgav on LTi-like ILC3s promote RORγt^+^ Treg homeostasis. **a**, Dot plot showing selected genes expressed in ILC3 (x axis) and Treg (y axis) cell clusters. Dots indicate known protein-protein interactions in the STRING database^73^, with dot sizes showing the sum of mean normalized expression in ILC3 clusters and Treg clusters in the scRNA-seq data. **b**, qPCR analysis of *Itgb1, Itgb3, Itgb5, Itgb6 and Itgb8* expression in sort-purified LTi-like ILC3s from LI-LP (n = 7) and mLN (n = 6), relative to *Hprt*. **c**, Sort-purified RORγt^+^CD4^+^ T cells and LTi-like ILC3s were co-cultured for 72 hours with blockades of indicated integrins (10 μg/mL), and frequency of RORγt^+^ Tregs were analyzed by flow cytometry. ND, not detected. **d**, Quantification of Itgav on LTi-like ILC3s and eTACs in mLN, and quantification of Itgav on ILC3s in LI-LP of *Itgav*^fl/fl^ and *Rorc*^cre^ x *Itgav*^fl/fl^ mice (n = 3-5). **e**, Representative flow cytometry plot of the frequency (% in CD4^+^ T cells) (left panel) and quantification (right panel) of RORγt^+^ Tregs and Th17 cells in mLN (top panel) and LI-LP (bottom panel) of *Itgav*^fl/fl^ and *Rorc*^cre^ x *Itgav*^fl/fl^ mice (n = 8-9). **f**, Representative flow cytometry plot of the frequency (left panel) and quantification (right panel) of RORγt^+^ Tregs (RORγt^+^FoxP3^+^ among Hh-specific CD4^+^ T cells) and Th17 cells (RORγt^+^FoxP3^-^ among Hh-specific CD4^+^ T cells) were analyzed in mLN (top panel) and LI-LP (bottom panel) of *Itgav*^fl/fl^ and *Rorc*^cre^ x *Itgav*^fl/fl^ mice (n = 4). Data in (**d**, **f**) are representative of two independent experiments. Data in e are pooled from two independent experiments. Data are shown as means ± SEM, statistics shown in (**c**) are obtained by one-way ANOVA with Tukey’s multiple comparisons test, statistics shown in (**e**, **f**) are obtained by unpaired Student’s *t*-test (two-tailed), ns, not significant; *p<0.05, **p<0.01, ****p<0.0001.

### ILC3-Treg interactions are altered in IBD

IBD is a human disease characterized by chronic inflammation of the GI tract with a substantial alteration of ILC3s^4,40–43^ and RORγt^+^ T lymphocytes^44–49^. However, a comprehensive definition of ILC3s and RORγt^+^ T cell interactions in human health and IBD is lacking. This provoked us to perform scRNA-seq on the ILC and T cell compartments from the inflamed tissue versus adjacent non-inflamed tissue of the intestine from a human IBD patient (**Extended Data Fig. 9a**).

Seventeen clusters were identified and visualized by UMAP (**Fig. 5a**). We separated the clusters of immune cells into T cells versus non-T cells as determined by expression of *CD3E* (**Fig. 5b**). Among non-T cells, cluster 4 was identified as ILC3s by expression of *ID2, KIT* and *RORC* (**Fig. 5c**), and representation of these cell types was reduced in the inflamed tissue relative to adjacent non-inflamed tissue (**Fig. 5d**). This decrease of ILC3s in the inflamed human intestine was associated with a significant decrease in expression of genes associated with ILC3 identity including *RORC, RORA, KIT, IL1R1, CCR6, IL22*, and those involved in antigen processing and presentation including *CD74, HLA-DRA, CD83, CD81, CTSH, IRF4* (**Fig. 5e**). Further, this impairment of ILC3s during intestinal inflammation was independently validated with colonic biopsies from a cohort of pediatric Crohn’s disease patients relative to age-matched controls (**Fig. 5f, Extended Data Fig. 9b**). Among the T cells, cluster 6 and 8 were identified as Tregs by expression of *CD4* and *FOXP3* (**Fig. 5g**). A direct comparison of differentially expressed genes revealed that cluster 6 represents RORγt^+^ Tregs as they displayed higher expression of *RORA, RORC, CCR6* and *CXCR6*, while cluster 8 exhibit features of thymic-derived Tregs such as higher expression of Helios/*IKZF2* (**Fig. 5h, Supplementary Table 2**). Representation of RORγt^+^ Tregs (cluster 6) was also reduced in the inflamed human intestine relative to matched non-inflamed tissue, and this was associated with a significant decrease in genes associated with RORγt^+^ Treg identity, function and TCR signaling (**Fig. 5i, j**). Several of these changes in abundance or transcriptional signatures among ILC3s and Tregs were found in another publicly available data set from human IBD samples^44^ (**Extended Data Fig. 10a-c**). Further, a significant reduction in the frequency of RORγt^+^ Tregs was also independently validated in colonic biopsies from a cohort of pediatric Crohn’s disease patients relative to age-matched controls (**Fig. 5k, Extended Data Fig. 9b, Supplementary Table 3**). Finally, there was a modest but significant positive correlation between the frequency of ILC3s and the frequency of RORγt^+^ Tregs in all analyzed intestinal biopsies while no striking correlation between the frequency of Th17 and RORγt^+^ Tregs was observed (**Fig. 5l, Extended Data Fig. 10d**). Importantly, similar results were validated in a second independent cohort of pediatric Crohn’s disease patients (**Extended Data Fig. 10e-g**). These data suggest that like our findings in mice, ILC3s promote RORγt^+^ Tregs to support intestinal health in humans, and this pathway becomes altered in IBD.

**Fig. 5.**
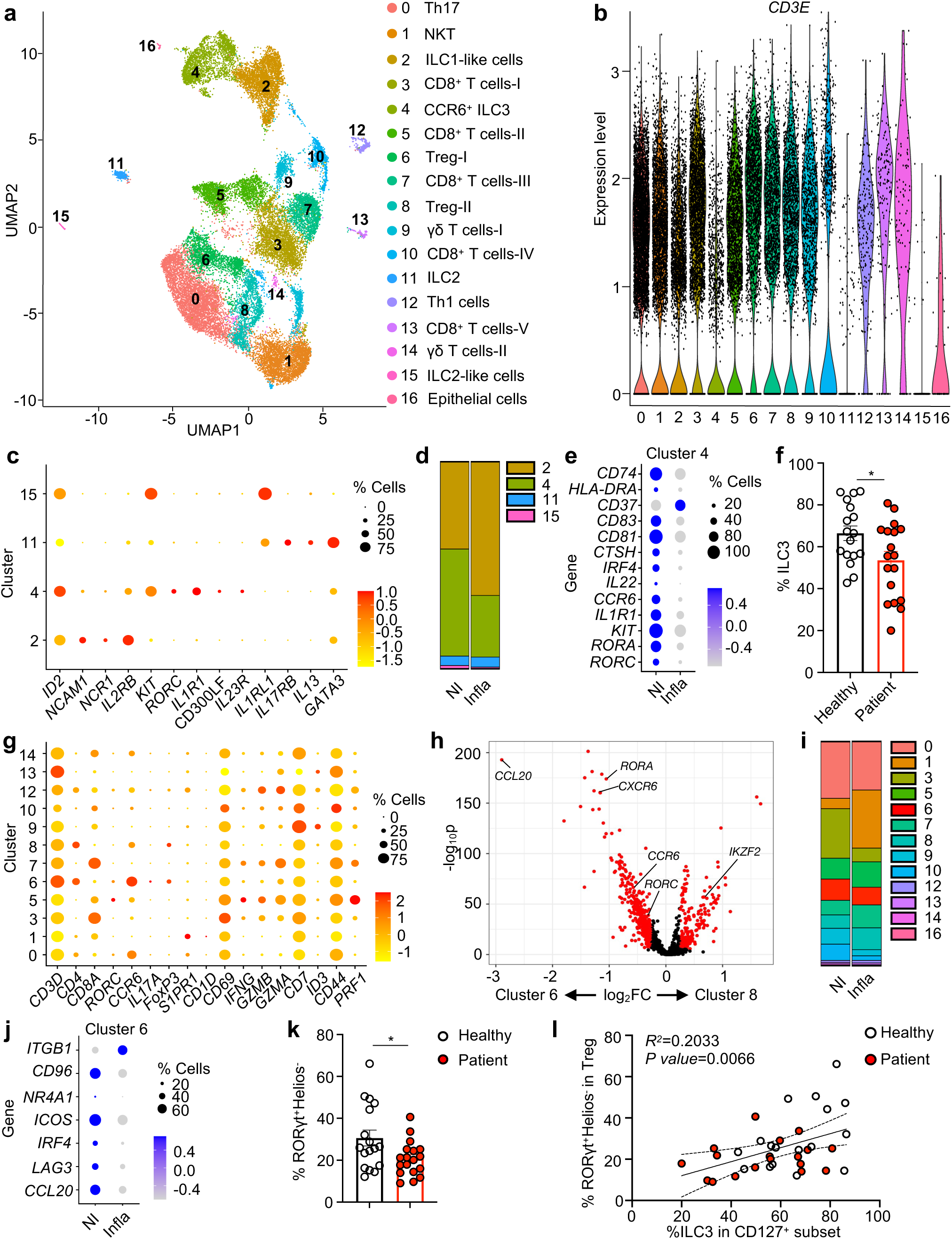
ILC3s and RORγt^+^ Tregs interactions are impaired in human IBD. **a**, UMAP plots of scRNA-seq data from ILCs and T lymphocytes in inflamed tissue and adjacent noninflamed tissue from a Crohn’s disease patient. ILCs and T lymphocytes were sort-purified for scRNA-seq, 13,072 cells from noninflamed tissue and 14,104 cells from inflamed tissue were sequenced. **b**, Violin plot showing the expression of *CD3E* among all the identified clusters. **c**, Dot plot showing the mean expression (color) of indicated genes in clusters grouped by low *CD3E* expression, dot size represents the proportion of cells in a cluster with the gene detected. **d**, Bar graph showing the composition of non-T lymphocytes in non-inflamed tissue (NI) versus inflamed tissue (Infla) from patient as indicated in (**a**). **e**, Expression of indicated genes in cluster 4 in non-inflamed versus inflamed tissue from patient as indicated in (**a**). **f**, Quantification of frequency of ILC3 in a cohort of IBD patients. Healthy donor n = 17, IBD patient n = 18. **g**, Dot plot showing the mean expression (color) of indicated genes in clusters grouped by high *CD3E* expression, dot size represents the proportion of cells in a cluster with the gene detected. **h**, Volcano plot of differentially expressed genes between cluster 6 and cluster 8 of scRNA-Seq data set from (**a**). **i**, Bar graph showing the composition of T lymphocytes across non-inflamed tissue versus inflamed tissue from patient as indicated in (**a**). **j**, Expression of indicated genes in cluster 6 in non-inflamed versus inflamed tissue from patient as indicated in (**a**). **k**, Quantification of frequency of RORγt^+^ Tregs among total Tregs in a cohort of IBD patients. Healthy donor n = 17, IBD patient n = 18. **l**, Correlation analyses between the ILC3 (ILC3 among CD127^+^ ILCs) and RORγt^+^ Tregs (RORγt^+^ Helios^-^ among FoxP3^+^ Tregs) in a cohort of healthy donor (n = 17) and IBD patients (n = 18). Data in (**f**, **k**) are shown as means ± SEM, statistics shown in (**f**, **k**) are performed using Mann–Whitney U-test (unpaired), correlative analyses in (**l**) are compared by Pearson’s rank correlation coefficient (*R^2^*), *p<0.05, **p<0.01, ****p<0.0001.

Our collective data sets indicate that LTi-like ILC3s critically select for RORγt^+^ Tregs and against Th17 cells to support immunologic tolerance to microbiota and maintain intestinal homeostasis (**Extended Data Fig. 10h**), which substantially broadens our current understanding of intestinal selection^26, 32^ and mucosal immunology. This is likely occurring after initial priming by DCs or other redundant antigen presenting cells in response to colonization with microbes, impacts T cells with distinct antigen-specificities, and requires antigen presentation via MHCII on LTi-like ILC3s with contributions from Itgav and gradients of competition for IL-2. The αvβ3 integrin could support processing of TGF-β or interact with other extracellular ligands that could be important in this process. Furthermore, our data sets demonstrate an unexpected interconverting potential between RORγt^+^ eTACs and LTi-like ILC3s. Therefore RORγt^+^ eTACs could represent a previously unappreciated progenitor, or simply a cell type that resides within lymph nodes that can interconvert to maintain the pool of MHCII^+^ ILC3s at distinct points of development or disease states. This is an area which requires additional investigation given a prior report on the enrichment of RORγt^+^ eTACs in early life lymph nodes and that this process is controlled RANK-RANKL interactions^18^. These are important findings in the overall emerging paradigm that there is sophisticated cross-regulation between RORγt-expressing lymphocytes that orchestrates intestinal health, which is essential to preserve the ability to rapidly respond to intestinal injury, infection, or inflammation. Our results indicate that LTi-like ILC3s play a non-redundant role in selecting for RORγt^+^ Tregs. Finally, we identify a positive correlation of ILC3s and RORγt^+^ Tregs in IBD patients and a fundamental disruption of these cell types in IBD. Intriguingly, each of these RORγt^+^ populations express IL-17, IL-23R and integrin *α*4*β*7, and it will be important to consider how targeting these pathways with currently available biologic therapies impacts their functional cellular interactions. Taken together, these findings fundamentally advance our understanding of the heterogeneity, lineage-relationships, and functional interactions by which RORγt expressing cells coordinate intestinal health and immune tolerance to the microbiota. Specifically, our data suggests that ILC3s, which are well-documented to become altered in human IBD, inherently underlies the dysregulation of microbiota-specific ROR*γ*t^+^ Tregs and Th17 cells, thus shifting the balance from tolerance to inflammation. Defining strategies to prevent this impairment of ILC3s, or promote their expansion, may hold the key to reinstating tolerance to microbiota in human IBD.

## Acknowledgements

We thank members of the Sonnenberg Laboratory for discussions and critical reading of the manuscript. Research in the Sonnenberg Laboratory is supported by the National Institutes of Health (R01AI143842, R01AI123368, R01AI145989, U01AI095608, R21CA249274, R01AI162936 and R01CA274534), an Investigators in the Pathogenesis of Infectious Disease Award from the Burroughs Wellcome Fund, the Meyer Cancer Center Collaborative Research Initiative, the Dalton Family Foundation, and Linda and Glenn Greenberg. Wenqing Zhou, J.G., L.Z. and Wen Zhang are supported by fellowships from the Crohn’s and Colitis Foundation (831404, 519428, 608975, and 901000, respectively). D.R.W. and F.G. are supported by a Senior Research Fellowship from the Wellcome Trust to DRW (110199/Z/15/Z). J.G.F. is supported by P30-ES002109 and R35CA210088. G.F.S. is a CRI Lloyd J. Old STAR. We would like to thank the Epigenomics Cores of Weill Cornell Medicine, Gregory Putzel, and all contributing members of the JRI IBD Live Cell Bank, which is supported by the JRI, Jill Roberts Center for IBD, Cure for IBD, the Rosanne H. Silbermann Foundation, the Sanders Family and Weill Cornell Medicine Division of Pediatric Gastroenterology and Nutrition. JRI Live Cell Bank consortium members include David Artis, Randy S. Longman, Gregory F. Sonnenberg, Ellen Scherl, Robbyn Sockolow, Dana Lukin, Robert Battat, Thomas Ciecierega, Aliza Solomon, Elaine Barfield, Kimberley Chien, Johanna Ferriera, Jasmin Williams, Shaira Khan, Peik Sean Chong, Samah Mozumder, Lance Chou, Wenqing Zhou, Anees Ahmed, Connie Zhong, Ann Joseph, Sanchita Kashyap, Joseph Gladstone, and Samantha Jensen.

## Author contributions

M.L., H.S. and G.F.S. conceived the project. M.L. performed most experiments and analyzed the data. L.K., F.G., J.G., Wenqing Zhou, L.Z., Wen Zhang and J.Z. helped with experiments and data analyses. J.G.F. Z.S., Y.F., T.M.L., G.E. and D.R.W. provided essential tools, scientific advice, and expertise. R.E.S. and JRI Live Cell Bank contributed to clinical sample acquisition and processing. M.L. and G.F.S. wrote the manuscript, with input from all the authors.

## Author information

H.S. is currently employed by EA Pharma Co., Ltd. Kanagawa, Japan. The authors declare no other competing interests. Correspondence and requests for materials should be addressed to.

## Materials and methods

### Mice

C57BL/6J mice, *Rag1^-/-^* mice, Thy1.1 transgenic mice, CD45.1 transgenic mice, Rosa-26- loxP-flanked STOP yellow fluorescent protein gene (eYFP) mice^50^, *H2-Ab1^fl/fl^* mice^51^, Red5^cre^ (*Il5*^cre^) mice^52^, *Foxp3*^cre^ mice^53^, *Cd4*^cre^ mice^54^, *Itgax*^cre^ (*Cd11c*^cre^) mice^55^, *Clec9a*^cre^ mice^56^, *Il22^cre^* mice^57^, *Rorc^fl/fl^* mice^58^, *Itgav^fl/fl^* mice^59^, SFB (7B8) TCR transgenic mice^33^ and *H. hepaticus* (HH7-2) TCR transgenic mice^34^ were purchased from the Jackson Laboratories. *Rag2*^-/-^*Il2rg*^-/-^ mice were purchased from Taconic Biosciences. *Rorc^cre^* mice and *Rorc(γt)*-*Gfp*^TG^ (RORγt-eGFP) mice^60^ were provided by Gerard Eberl. *Aire^cre^* mice^61^ were provided by Yong Fan. MHCII^ΔILC3^ mice were generated as previously described^32^. MHCII^ILC3+^ mice were generated by crossing *Rorc*^cre^ mice with IAb^b^STOP^fl/fl^ mice^62^ provided by Terri M. Laufer. *Ncr1*^cre^ mice^63^ were kindly provided by Eric Vivier (Inserm). *Il7r*^cre^ mice^64^ were kindly provided by David Artis (Weill Cornell Medicine) with permission from H.R. Rodewald. *Il2^fl/fl^* mice^65^ were kindly provided by Kendall Smith (Weill Cornell Medicine). All mice were on a C57BL/6 background and maintained in specific pathogen-free facilities in Weill Cornell Medicine. Male and female mice were used at 6 to 12 weeks of age unless otherwise indicated. All protocols were approved by the Institutional Animal Care and Use Committee at Weill Cornell Medicine, and all experiments were performed in accordance with its guidelines.

### Flow cytometry and cell sorting

Single cell suspensions were incubated on ice with conjugated antibodies in PBS containing 2% FBS and 1 mM EDTA. Unlabeled anti-CD16/32 (clone 2.4G2, BD biosciences) was used to block Fc receptors when analyzing myeloid cells. Dead cells were excluded with Fixable Aqua Dead Cell Stain (Thermo Fisher Scientific). The staining antibodies for flow cytometry were mainly purchased from Thermo Fisher Scientific, Biolegend or BD Biosciences. For mouse cell-surface staining: CCR6 (29-2L17), NKp46 (29A1.4), CD3ε (145-2C11), CD4 (RM4-5, GK1.5), CD5 (53-7.3), CD8α (53-6.7), CD11b (M1/70), CD11c (N418), CD19 (1D3), Gr1 (RB6-8C5), Ly-6G (1A8), CD45R/B220 (RA3-6B2), CD45.1 (A20), CD45.2 (104), CD45 (30-F11), CD64 (X54-5/7.1), CD90.1 (OX-7), CD90.2 (30-H12), CD127 (A7R34), CD51 (RMV-7), F4/80 (BM8), FcεR1α (MAR-1), MHC-II (M5/114.15.2), NK1.1 (PK136), TCRγδ (GL3), KLRG1 (2F1/KLRG1), CD44 (IM7), CD62L (MEL-14), CD25 (PC61), CXCR6 (SA051D1), Siglec-G (SH1), CD26 (H194-112), Nur77 (12.14), CD172α (P84), XCR1 (ZET), Ly-6C (HK1.4), CD132 (TUGm2), CD138 (281-2), CD117 (ACK2), CD122 (TM-β1), Sca-1 (D7), and Integrin α4β7 (DATK32). For mouse intracellular staining: FoxP3 (FJK-16S), GATA3 (L50-823), IL-17A (17B7), IFN-γ (XMG1.2), Ki-67 (SolA15), Bim (C34C5), RORγt (B2D) and T-bet (4B10). Lineage markers for mouse are: CD3ε, CD5, CD19, B220, Gr1, NK1.1, CD11b, CD11c unless otherwise indicated. For human samples cell-surface staining: CD3 (UCHT1), CD11c (3.9), CD14 (TuK4), CD19 (HIB19), CD34 (581), CD4 (SK3), CD45 (HI30), CD25 (BC96), CD94 (DX22), CD117 (104D2), CD123 (6H6), CD127 (A019D5), FcεR1 (AER-37(CRA1)) and NKp44 (44.189). For human samples intracellular staining: FOXP3 (PCH101), HELIOS (22F6). Human RORγt^+^ Tregs were gated as CD45^+^CD3^+^CD4^+^FOXP3^+^HELIOS^-^RORγt^+^, human ILC3s were gated as CD45^+^CD3^-^ CD11c^-^CD14^-^CD19^-^CD34^-^CD94^-^CD123^-^FcεR1^-^CD127^+^CD117^+^.

For intracellular staining, cells were fixed and permeabilized with FoxP3/Transcription Factor Staining Buffer Set following manufacture’s instruction (Thermo Fisher Scientific). Briefly, cells were incubated with Foxp3 Fixation/Permeabilization working solution for 30 min at room temperature or overnight at 4°C, then stained with intracellular targets by incubating with following conjugated antibodies in 1X Permeabilization Buffer for 30 min at room temperature. For intracellular cytokine staining, cells were first incubated for 4 h in RPMI with 10% FBS, 50 ng/ml phorbol 12-myristate 13-acetate (PMA), 750 ng/ml ionomycin and 10 μg/ml brefeldin A, all obtained from Sigma Aldrich. Antibodies for flow cytometry were purchased from BioLegend, Thermo Fisher Scientific or BD Biosciences. IL-2 binding capacity was assessed using a biotinylated IL-2 fluorokine assay kit (R&D Systems), following the manufacturer’s instructions. Flow cytometry data collection was performed with an LSR Fortessa (BD Biosciences) and analyzed with FlowJo V10 software (Tree Star, Inc). Cell sorting was performed with an Aria II (BD Biosciences).

### Quantitative PCR

Sort-purified cells were lysed in RLT buffer (Qiagen). RNA was extracted via RNeasy mini kits (Qiagen), as per the manufacturer’s instructions. Reverse transcription of RNA was performed using Superscript reverse transcription according to the protocol provided by the manufacturer (Thermo Fisher Scientific). Real-time PCR was performed on cDNA using SYBR green chemistry (Applied Biosystems). Reactions were run on a real-time PCR system (ABI 7500; Applied Biosystems). Samples were normalized to *Hprt1* and displayed as a fold change compared to controls.

### Preparation of single cell suspensions from intestine or lymph nodes

Small intestine, cecum and colon were removed, opened longitudinally, and rinsed with ice-cold PBS. Peyer’s patches were removed from small intestine. Dissected intestinal tissues were cut into approximately 1.5 cm pieces and intestinal epithelial cells were dissociated by incubating in HBSS (Sigma-Aldrich) containing 5 mM EDTA (Thermo Fisher Scientific), 1 mM DTT (Sigma-Aldrich) and 2% FBS with shaking at 200 rpm for 20 min at 37°C. Epithelial cells dissociation was performed twice. Samples were vortexed and rinsed with PBS after each step. Epithelial cells fraction was discarded. Remaining tissues were enzymatically digested in RPMI containing 0.4 U/mL dispase (Thermo Fisher Scientific), 1 mg/mL collagenase III (Worthington), 20 μg/mL DNase I (Sigma-Aldrich) and 10% FBS on a shaker for 45 min at 37°C. Leukocytes were enriched by Percoll gradient centrifugation (GE-Healthcare). Mesenteric lymph nodes were chopped and incubated in RPMI containing 2% FBS and 2 mg/mL Collagenase D (Sigma-Aldrich) with shaking at 200 rpm for 20 min at 37°C, cells were then dissociated using a pasteur pipette, and filtered through 70 μm cell strainer in PBS containing 0.5 mM EDTA and 2% FBS.

### Adoptive transfer of microbiota specific TCR transgenic T cells

Mice (CD45.2^+^, CD90.2^+^) were colonized with *H. Hepaticus* (ATCC 51449/Hh3B1) by oral gavage 7 days before T cell transfer as described previously^34^, in some experiments antibiotics (cocktail containing 1 g/L ampicillin, 1 g/L colistin, 5 g/L streptomycin in drinking-water for 3 days) were applied to help with a better colonization of *H. Hepaticus*. SFB is continuously colonizing mice in our facility. Naïve CD4^+^ T cells of Hh7-2 (CD45.1^+^, CD90.2^+^) and SFB7B8 (CD45.2^+^, CD90.1^+^) mice were isolated from spleen and lymph nodes by a naïve CD4^+^ T cell isolation kit following manufacturer’s instructions (Miltenyi Biotech) or by FACSAria cell sorter (BD Biosciences) gated as CD45^+^CD3^+^CD4^+^CD25^−^CD44^low^CD62L^high^. Recipient mice received 10,000-100,000 cells/mouse of both Hh7-2 and SFB7B8 CD4^+^ T cells retro-orbitally and were analyzed 2 weeks after CD4^+^ T cell transfer.

### Adoptive transfer of sort purified ILC3s and RORγt^+^ eTACs to Rag2/Il2rg double knockout mice

RORγt-eGFP reporter mice or *Aire*^cre^ x Rosa26^lsl-YFP^ fate mapped mice were used as donor mice, LTi-like ILC3s (gated as CD45^+^CD3^-^CD5^-^CD19^-^B220^-^Gr1^-^NK1.1^-^CD11b^-^CD11c^-^KLRG1^-^ GFP^+^CCR6^+^CD127^+^CD90.2^+^), RORγt^+^ eTACs (gated as CD45^+^CD3^-^CD5^-^CD19^-^B220^-^Gr1^-^NK1.1^-^ CD11b^-^CD11c^-^KLRG1^-^GFP^+^CCR6^+^CD127^-^CD90.2^-^SiglecG^+^DPP4^+^) or SiglecG^+^Dpp4^+^YFP^+^ cells were sort-purified from mLN by FACSAria cell sorter (BD Biosciences). Recipient mice received 7,000 ILC3s, 1,800 eTACs or 8,500 YFP^+^ cells retro-orbitally and were analyzed 6-8 weeks after adoptive cell transfer.

### Immunofluorescence and image analysis

Tissue sections from experimental mice were cut and stained as described previously^66, 67^. Briefly, 6-μm-thick sections of tissue were cut, fixed in cold acetone at 4°C for 20 min and then stored at −20°C before staining. The detection of RORγt in frozen tissue sections using immunofluorescence has been described previously^68^. Antibodies raised against the following mouse antigens were used: purified Armenian hamster anti-mouse CD3 (clone 145-2C11, 1:100, Biolegend), CD11c (Clone HL3 - Isotype, 1:100, BD Pharmingen™), rat anti-mouse IL-7Rα eFluor660 (clone A7R34, 1:25, Thermo Fisher), biotin anti-mouse FOXP3 (clone: FJK-16s, 1:50, Thermo Fisher) and rat anti-mouse RORγt (clone AFKJS-9, 1:25, Thermo Fisher). Detection of RORγt expression required amplification of the signal as described previously^68^. Purified RORγt antibodies were detected with donkey anti-rat-IgG-FITC (1:150, Jackson ImmunoResearch), and then rabbit anti-FITC-AF488 (1:200, Invitrogen) and then with donkey anti-rabbit-IgG-AF488 (1:200, Invitrogen). Purified anti-CD3 antibodies were detected with DyLight™ 594 goat anti-Armenian hamster IgG (clone Poly4055, 1:200, Biolegend) and biotinylated FOXP3 antibodies were detected with SA-AF555 (1:500, Invitrogen). Sections were counterstained with DAPI (Invitrogen) and mounted using ProLong Gold (Invitrogen). Slides were analyzed on a Zeiss 780 Zen microscope (Zeiss). High resolution (x63) pictures of the interfollicular areas were taken. The number of RORγt^+^ Tregs was enumerated using the Zen software (Zeiss). This was performed by examining 8 to 12 images of interfollicular zones from at least 3 individual mice. For each ROR*γ*t^+^ Treg in the image was then quantified as either ‘adjacent’ or ‘not adjacent’ to an ILC3 based on observed co-localization of markers on the surface membrane (CD3 for T cell membrane and IL-7R but not CD3 for ILC3 membrane in this context, in addition to intracellular transcription factors).

### In vitro ILC3 and T cell culture

T cell stability assay was conducted as previously reported ^69^ with minor modifications. Sort-purified RORγt^+^ CD4^+^ T cells (CD45^+^CD5^+^CD4^+^RORγt^GFP+^) were plated in a round-bottom 96-well plate with 1:1 or 2:1 ratio (5,000 to 10,000 cells/well) of ILC3s (CD45^+^CD5^-^B220^-^TCRγδ^-^KLRG1^-^NK1.1^-^CD11c^-^RORγt^GFP+^). Large intestine and mLN were used to collect RORγt^+^ CD4^+^ T cells and ILC3s. Cells were incubated in DMEM with high glucose, supplemented with 10% FBS, 1 mM Sodium Pyruvate, 10 mM HEPES, non-essential amino acid, 80 μM 2-mercaptoethanol, 100 U/mL penicillin, 100 μg/mL streptomycin (all from Thermo Fisher Scientific), 10 ng/mL recombinant IL-7 (Thermo Fisher Scientific), anti-integrin αV (Biolegend), anti-integrin β1 (BD Biosciences), anti-integrin β3 (BD Biosciences) and anti-integrin αVβ6 (MilliporeSigma) at 10 μg/mL as indicated at 37°C/5% CO_2_ for 72 hours. Cells were analyzed by flow cytometry.

### Single cell RNA-Sequencing

CD45^+^TCRβ^+^GFP^+^ (T cells) and CD45^+^TCRβ^-^GFP^+^ (non-T cells) were sorted from the mLN of RORγt-eGFP mice and mixed equally at 1:1 ratio to enrich the non-T cell populations. CD45^+^CD19^-^CD14^-^CD123^-^FcεRIα^-^CD34^-^CD94^-^CD4^-^CD127^+^ILCs and CD45^+^CD3^+^ T cells were sorted from the resection of inflamed Crohn’s disease patient versus adjacent tissue and mixed at 1:4 ratio. scRNA-Seq libraries were generated using the 10X Genomics Chromium system with 3’ version 3 chemistry. Libraries were sequenced on an Illumina NovaSeq instrument. Reads were processed by 10X’s Cell Ranger version 3.1.0 using the mm10 reference genome, resulting in a filtered HDF5 file. scRNA-Seq data were further processed and analyzed using R version 3.6.3 (R Core Team 2020) and the Seurat package version 3.2.3^70^. Specifically, Cell Ranger output was imported using the Read10X_h5 function. Seurat objects were created using only genes appearing in at least 3 cells. Cells were further filtered to exclude those with fewer than 600 genes detected, more than 5000 genes detected, or more than 10 percent mitochondrial reads. Read counts were then normalized using the NormalizeData function. The graph representing cells with similar expression patterns was generated with the FindNeighbors function using the 20 largest principal components. Cell clusters were generated using the Louvain algorithm implemented by the FindClusters function with resolution parameter equal to 0.4. Marker genes for each cluster were determined using the Wilcoxon test on the raw counts, implemented by the function FindAllMarkers, and including only positive marker genes with log fold changes greater than 0.25 and Bonferroni-corrected *p* values less than 0.01. Cluster names were determined by manual inspection of the lists of cluster marker genes. Dimensionality reduction by Uniform Manifold Approximation and Projection was performed using the RunUMAP function with the 20 largest principal components. All visualizations of scRNA-Seq data were generated using the Seurat package as well as ggplot2 version 3.3.3^71^.

### Trajectory analysis

Principal components analysis (PCA) as well as dimensionality reduction by UMAP, were performed using Seurat on the subset of cells belonging to non-T cell clusters (2,6, and 11). The resulting Seurat object was converted for use in Slingshot using the as SingleCellExperiment of function of Seurat. Pseudotime analysis was performed using version 2.0.0 of the Slingshot R package^72^.

### Cell-cell interaction network

Selected genes expressed in ILC3 clusters (2 and 12) as well as selected genes expressed in Treg clusters (0, 5, and 13) were submitted to the STRING database^73^. The resulting list of known protein-protein interactions was filtered to include only entries with either database annotations (“database_annotated”) or experimental evidence (“experimentally_determined_in-teraction”).

### Human intestinal tissue isolation

De-identified intestinal biopsies from the colon of pediatric individuals with Crohn’s disease and sex- and age-matched controls with no IBD were obtained following Institutional Review Board approved protocols from the JRI IBD Live Cell Bank Consortium at Weill Cornell Medicine. Informed consent was obtained from all subjects. Biopsies were cryopreserved in 90% FBS and 10% DMSO for future side-by-side comparison. Following thawing, tissues were incubated in 0.5 mg/ml collagenase D and 20 mg/ml DNase I for 1 hour at 37°C with shaking. After digestion, remaining tissues were further dissociated mechanically by a syringe plugger. Cells were filtered through a 70 mm cell strainer and used directly for staining.

### Statistics

P values of datasets were determined by unpaired two-tailed Student’s *t*-test with 95% confidence interval for comparison between two independent groups. For repeated comparison between two groups, multiple unpaired two-tailed *t*-test was used. Where appropriate, Mann– Whitney U-test was performed. One-way ANOVA followed by Turkey’s multiple comparison test was used for comparison between more than two groups. Simple linear regression was used to analyze correlation. All statistics were performed with GraphPad Prism version 9 (GraphPad Software Inc.). P values of less than 0.05 were considered to be significant. *p<0.05; **p<0.01, ***p<0.001, ****p<0.0001 and ns, not significant. Investigators were not blinded to group allocation during experiments.

### Data availability

All data necessary to understand and evaluate the conclusions of this paper are provided in the manuscript and supplementary materials. scRNA-Seq data can be found through Gene Expression Omnibus under accession number GSExxxxxx.

---

**Extended Data Fig. 1.**
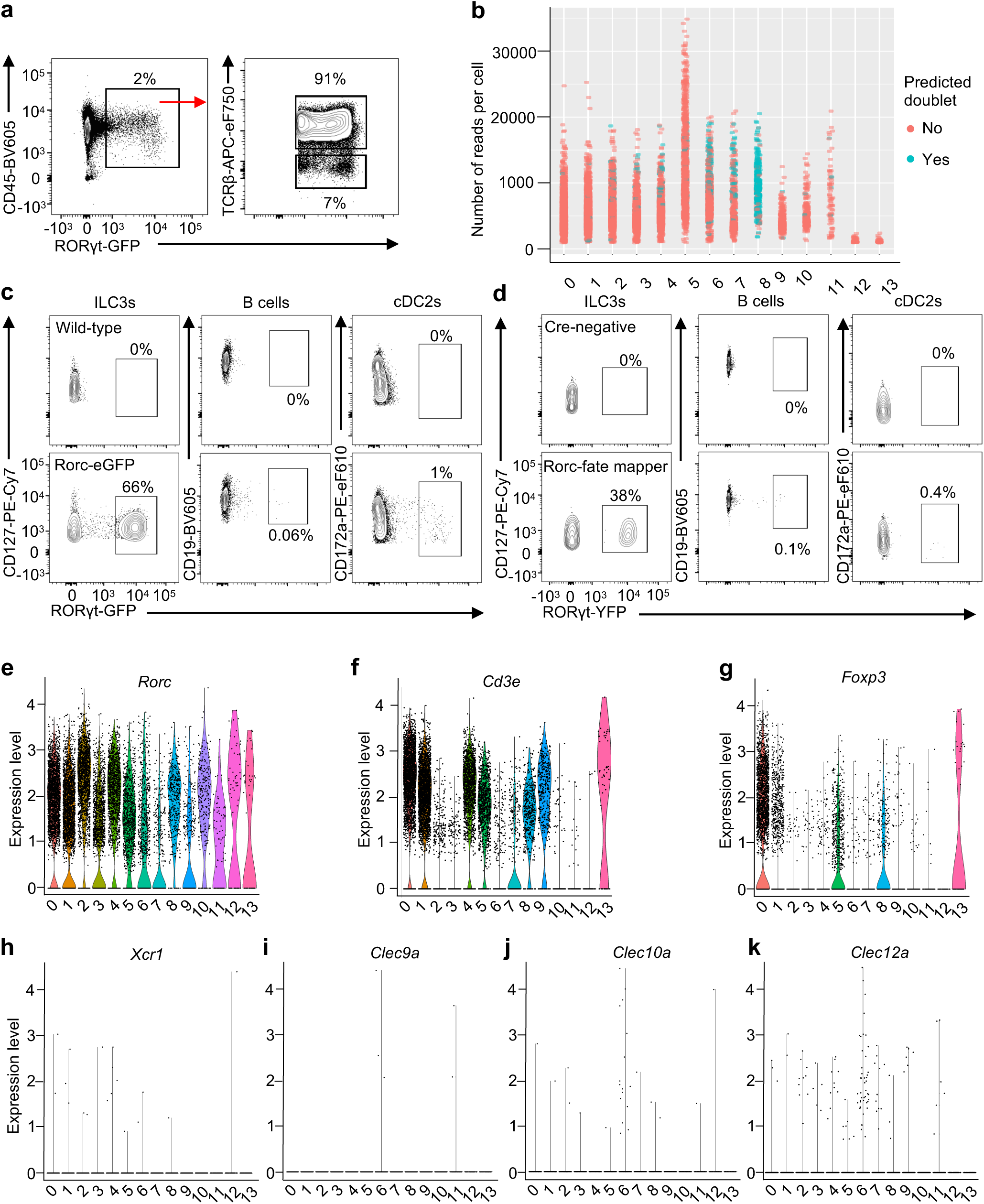
scRNA-seq profiling of RORγt^+^ cells from mouse mLN. **a**, Gating strategy to sort GFP^+^TCRβ^+^ and GFP^+^TCRβ^-^ cells (1:1, n = 3) for scRNA-seq. **b**, Doublet test showing cluster 8 as doublets. **c**, **d**, Representative flow cytometry plot of the frequency of RORγt^+^ cells in CD127^+^ ILC fractions (CD45^+^CD3ε^-^CD5^-^NK1.1^-^Ly6G^-^TCRγ/δ^-^B220^-^CD11b^-^CD11c^-^ KLRG1^-^CD127^+^), CD19^+^ B cell fractions (CD45^+^CD19^+^) and CD172a^+^ cDC2 fractions (CD45^+^ CD3ε^-^CD5^-^NK1.1^-^Ly6G^-^TCRγ/δ^-^B220^-^CD64^-^CD11c^+^MHCII^+^XCR1^-^CD172a^+^) from RORγt-eGFP reporter mice (n = 3) (**c**) and *Rorc*^cre^ x Rosa26^lsl-YFP^ fate mapped mice (n = 3) (**d**). **e**-**k**, Violin plot showing the expression of *Rorc* (**e**)*, Cd3e* (**f**)*, Foxp3* (**g**)*, Xcr1* (**h**)*, Clec9a* (**i**)*, Clec10a* (**j**)*, Clec12a* (**k**) among all the identified clusters.

**Extended Data Fig. 2.**
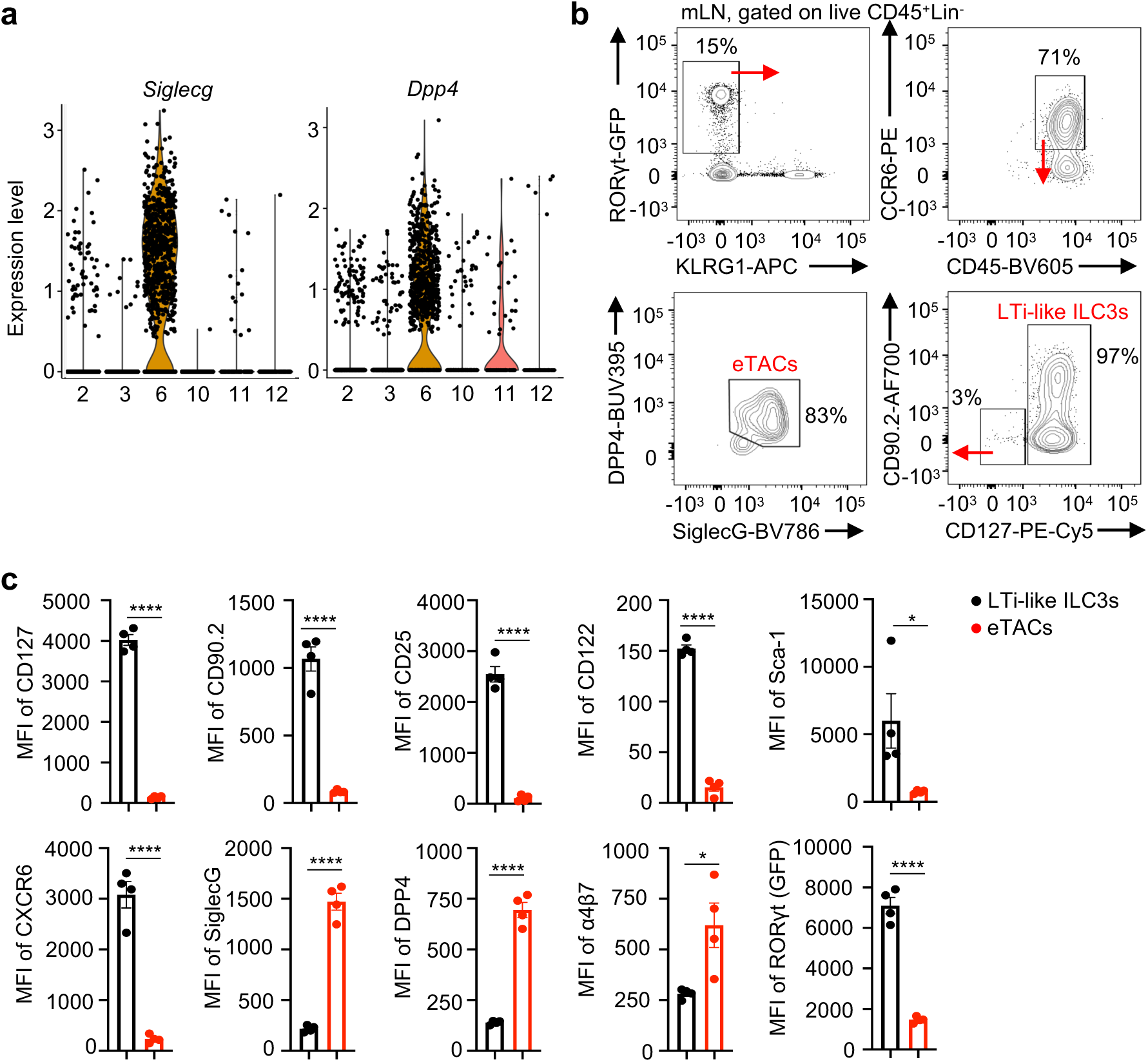
Characterization of RORγt^+^ eTACs and LTi-like ILC3s in mouse mLN. **a**, Violin plot showing the expression of *Siglecg* and *Dpp4* among all the identified clusters of non-T lymphocytes. **b**, Gating strategy to identify ILC3s and RORγt^+^ eTACs from mLN of RORγt-eGFP reporter mice (n = 4) for data shown in Fig. 1e, f. **c**, Quantification of indicated genes expression in LTi-like cells and RORγt^+^ eTACs shown in Fig. 1e (n = 4). Data in **c** are representative of three independent experiments. Data are shown as means ± SEM, statistics shown in (**c**) are obtained by unpaired Student’s *t*-test (two-tailed), ns, not significant; *p<0.05, **p<0.01, ***p<0.001, ****p<0.0001.

**Extended Data Fig. 3.**
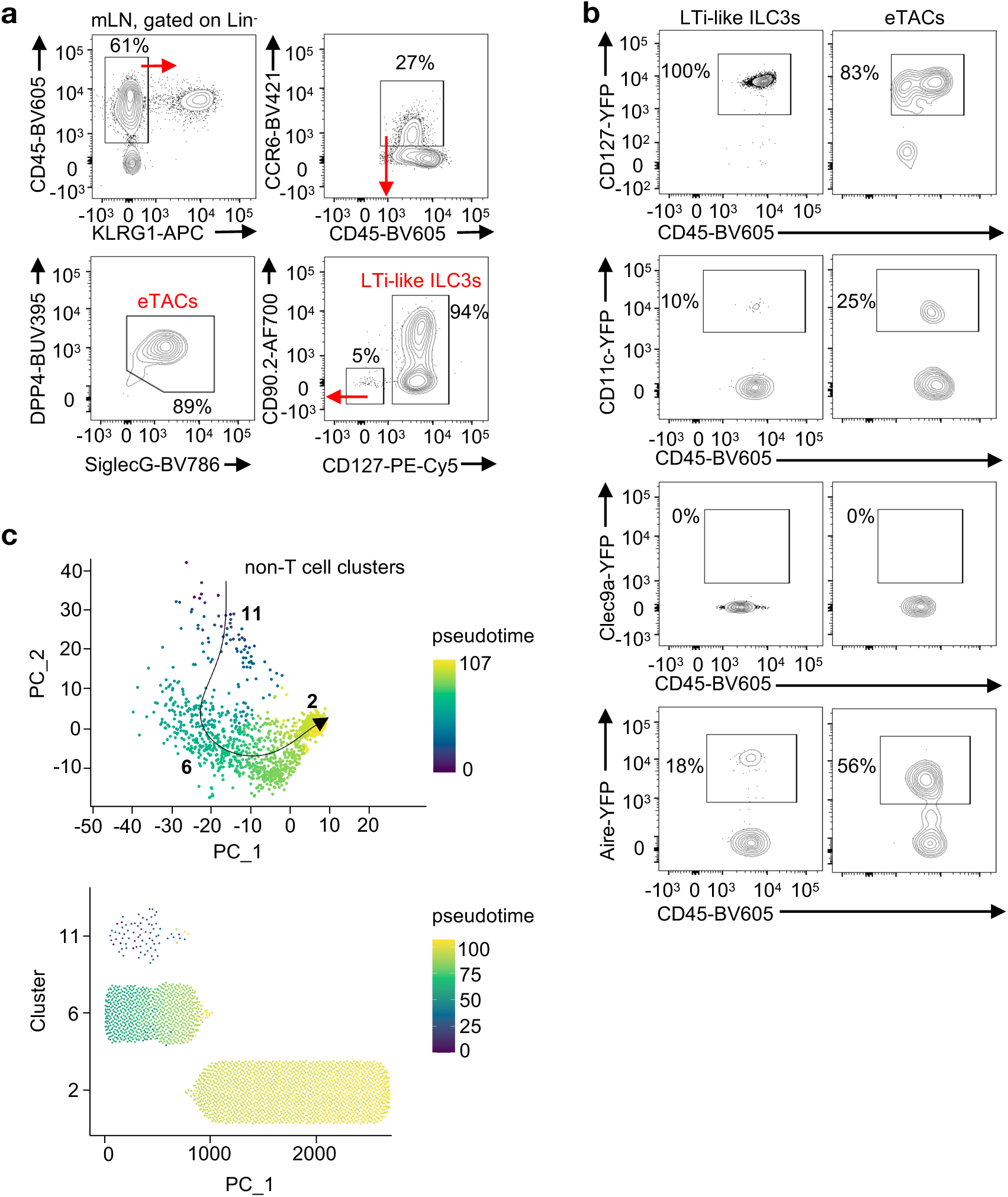
Fate mapping and trajectory analyses of RORγt^+^ eTACs and LTi-like ILC3s in mouse mLN. **a**, Gating strategy to identify LTi-like ILC3s and RORγt^+^ eTACs for fate mapping analyses in Fig. 1g, h. **b**, Representative flow cytometry plots showing expression of CD127, CD11c, Clec9a and Aire among “fate-mapped” LTi-like ILC3s and RORγt^+^ eTACs in mLN shown in Fig. 1g, h (n = 3-4). **c**, Potential developmental trajectory of indicated clusters inferred by pseudotime analysis plotted on PCA.

**Extended Data Fig. 4.**
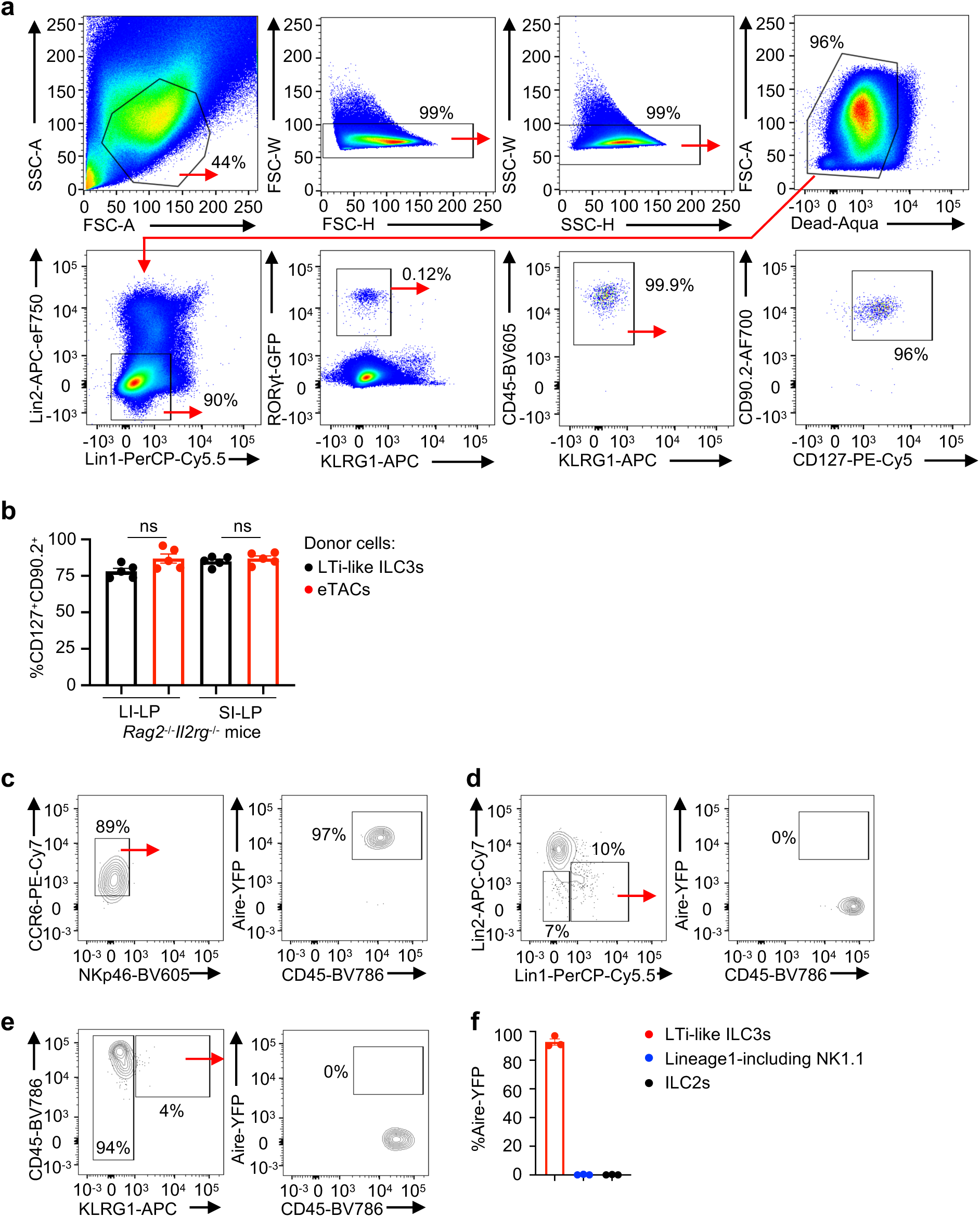
Gating strategy for ILC3 and fate mapping experiments. **a**, Gating strategy to show CD127^+^CD90.2^+^ ILC3s in small and large intestines of *Rag2^-/-^Il2rg^-/-^* recipient mice (n = 5) with adoptive transferred LTi-like ILC3 and RORγt^+^ eTACs sort purified from RORγt-eGFP reporter mice shown in Fig. 1j, k. **b**, Frequencies of CD127^+^CD90.2^+^ ILC3s in large and small intestine of *Rag2^-/-^Il2rg^-/-^* recipient mice (n = 5) with adoptive transferred LTi-like ILC3 and RORγt^+^ eTACs sort purified from RORγt-eGFP reporter mice. **c**-**f**, Representative flow cytometry plots of frequency (**c**, **d**, **e**) and quantification (**f**) of expression of Aire among “fate-mapped” LTi-like ILC3s, NK1.1^+^ cell subset and ILC2s in small intestine of *Rag2^-/-^Il2rg^-/-^* recipient mice (n = 3) with adoptive transferred siglecG^+^Dpp4^+^YFP^+^ cells sort purified from *Aire*^cre^ x Rosa26^lsl-YFP^ fate mapped mice. Data in (**b**-**f**) are representative of two independent experiments, shown as mean ± SEM, statistics shown in (**b**) are obtained by multiple unpaired Student’s *t*-test, ns, not significant.

**Extended Data Fig. 5.**
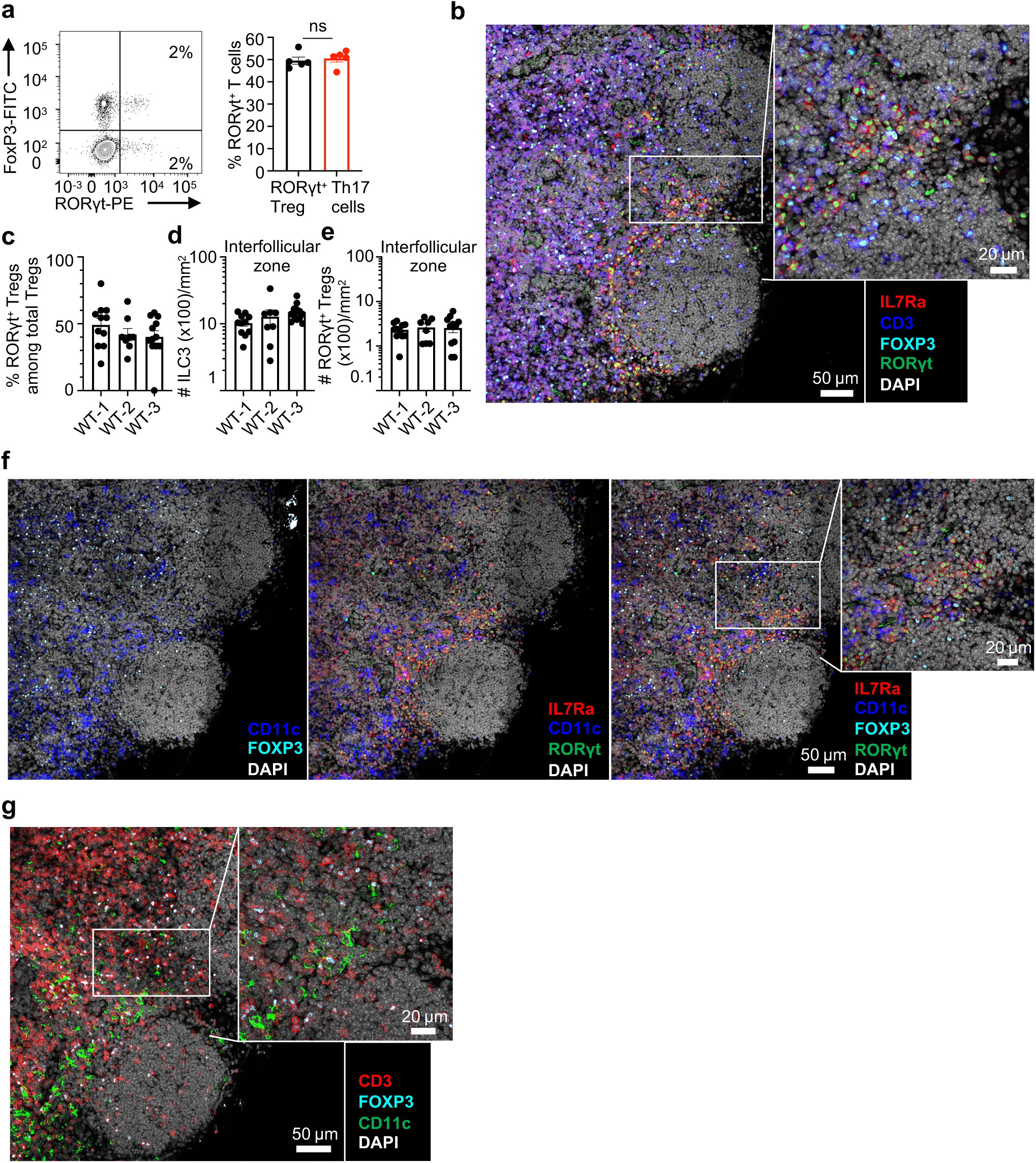
Immunofluorescence and quantification of different cell types in mLN. **a**, Representative flow cytometry plots of frequency (left) and percentage (right) of RORγt^+^FoxP3^+^ Tregs and RORγt^+^FoxP3^-^ Th17 cells among total RORγt^+^CD4^+^ T cells in mLN of WT mice (n = 5). **b**, Tile-scanned (left) and magnified (right) images of mLN stained for expression of IL7Rα (red), CD3 (blue), FOXP3 (cyan), RORγt (green) and DAPI (grey). **c-e**, Quantification of percentage of RORγt^+^Tregs among total Tregs (**c**), total numbers per mm^2^ of ILC3s (**d**) and RORγt^+^Tregs (**e**) in interfollicular zone of mLN of WT mice (n = 3). **f**, Tile-scanned images and serial sections of mLN stained for expression of IL7Rα (red), CD11c (blue), FOXP3 (cyan), RORγt (green) and DAPI (grey). Left panel is without IL7Rα and RORγt staining, middle panel is without FOXP3 staining, and right panel is a merge with a magnified image. **g**, Tile-scanned (left) and magnified (right) images of mLN stained for expression of CD3 (red), FOXP3 (cyan), CD11c (green) and DAPI (grey). Scale bar: 50 μm, 20 μm (in magnified images).

**Extended Data Fig. 6.**
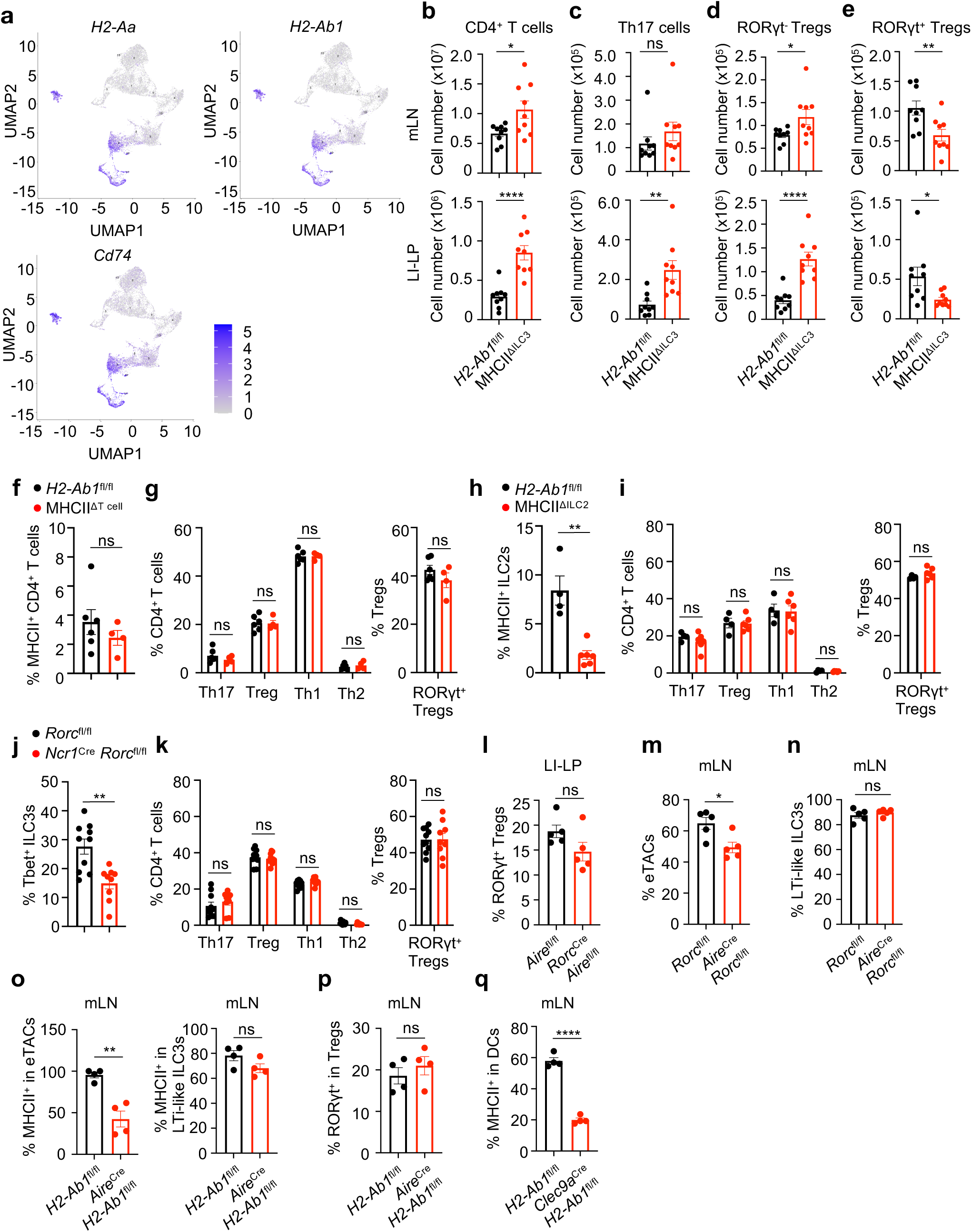
MHCII^+^ ILC3s selectively regulates T cell homeostasis in the gut. **a**, UMAP plots of scRNA-seq data showing expression of *H2-Ab1*, *H2-Ab1* and *Cd74* are enriched in clusters of LTi-like ILC3s and RORγt^+^ eTACs across all the identified clusters in mouse mLN. **b**-**e,** Cell numbers of each subset of CD4^+^ T cells in mesenteric lymph node (mLN, upper panel) and large intestine (LI, lower panel) of *H2-Ab1*^fl/fl^ and MHCII^ΔILC3^ mice (n = 8, pooled from 2 independent experiments). Total CD4^+^ T cells (**b**, CD45^+^CD3^+^CD4^+^); Th17 cells (**c**, CD45^+^CD3^+^CD4^+^ Foxp3^-^RORγt^+^); RORγt^-^Tregs (**d**, CD45^+^CD3^+^CD4^+^Foxp3^+^RORγt^-^); RORγt^+^Tregs (**e**, CD45^+^CD3^+^CD4^+^ Foxp3^+^RORγt^+^). **f**, **g**, Large intestine of *H2-Ab1*^fl/fl^ and *Cd4*^cre^ *H2-Ab1*^fl/fl^ (MHCII^ΔT cell^) mice were analyzed. Proportion of MHCII-expressing CD4^+^ T cells (**f**). Frequency of each subset among CD4^+^ T cells and RORγt^+^Tregs among total Tregs (**g**) (n = 4-6). **h**, **i**, Large intestine *of H2-Ab1*^fl/fl^ and *Red5*^cre^ *H2-Ab1*^fl/fl^ (MHCII^ΔILC2^) mice were analyzed. Proportion of MHCII-expressing ILC2s (**h**). Frequency of each subset among CD4^+^ T cells and RORγt^+^Tregs among total Tregs (**i**) (n = 4-6). **j**, **k**, Large intestine *of Rorc*^fl/fl^ and *Ncr1*^cre^ *Rorc*^fl/fl^ mice were analyzed. Proportion of T-bet-expressing ILC3s (**j**). Frequency of each subset among CD4^+^ T cells and RORγt^+^Tregs among total Tregs (**k**) (n = 9-10). Th17: Foxp3^-^RORγt^+^; Treg: Foxp3^+^; Th1: Foxp3^-^RORγ^-^T-bet^+^; Th2: Foxp3^-^RORγt^-^Gata3^+^. Frequency of each subset of CD4^+^ T cells was analyzed among total CD4^+^ T cells and frequency of RORγt^+^ Tregs was analyzed among Tregs. **l**, Quantification of RORγt^+^ Tregs among total CD4^+^ T cells in LI-LP of *Aire*^fl/fl^ and *Rorc*^cre^ x *Aire*^fl/fl^ mice (n = 5). **m**, **n**, Quantification of eTACs among total CD127^-^CD90^-^ cells (**m**) and LTi-like ILC3s among CD45^+^CCR6^+^ cells (**n**) in mLN of *Rorc*^fl/fl^ and *Aire*^cre^ x *Rorc*^fl/fl^ mice (n = 5). **o**, **p**, Quantification of MHCII expression among eTACs (**o,** left), LTi-like ILC3s (**o,** right) and RORγt^+^ Tregs among total Tregs (**p**) in mLN of *H2-Ab1*^fl/fl^ and *Aire*^cre^ x *H2-Ab1*^fl/fl^ mice (n = 4). **q**, Quantification of MHCII expression among DCs in mLN of *H2-Ab1*^fl/fl^ and *Clec9a*^cre^ x *H2-Ab1*^fl/fl^ mice (n = 4). Data are representative of two independent experiments unless otherwise indicated. Data shown as mean ± SEM. Statistics in (**f**, **h**, **j**, **l-q,** right of **g, i, k**) are obtained by two-tailed unpaired Student’s *t*-test. Statistics shown in (left of **g**, **i**, **k**) are obtained by multiple unpaired *t*-test. ns, not significant; *p<0.05, **p<0.01, ****p<0.0001.

**Extended Data Fig. 7.**
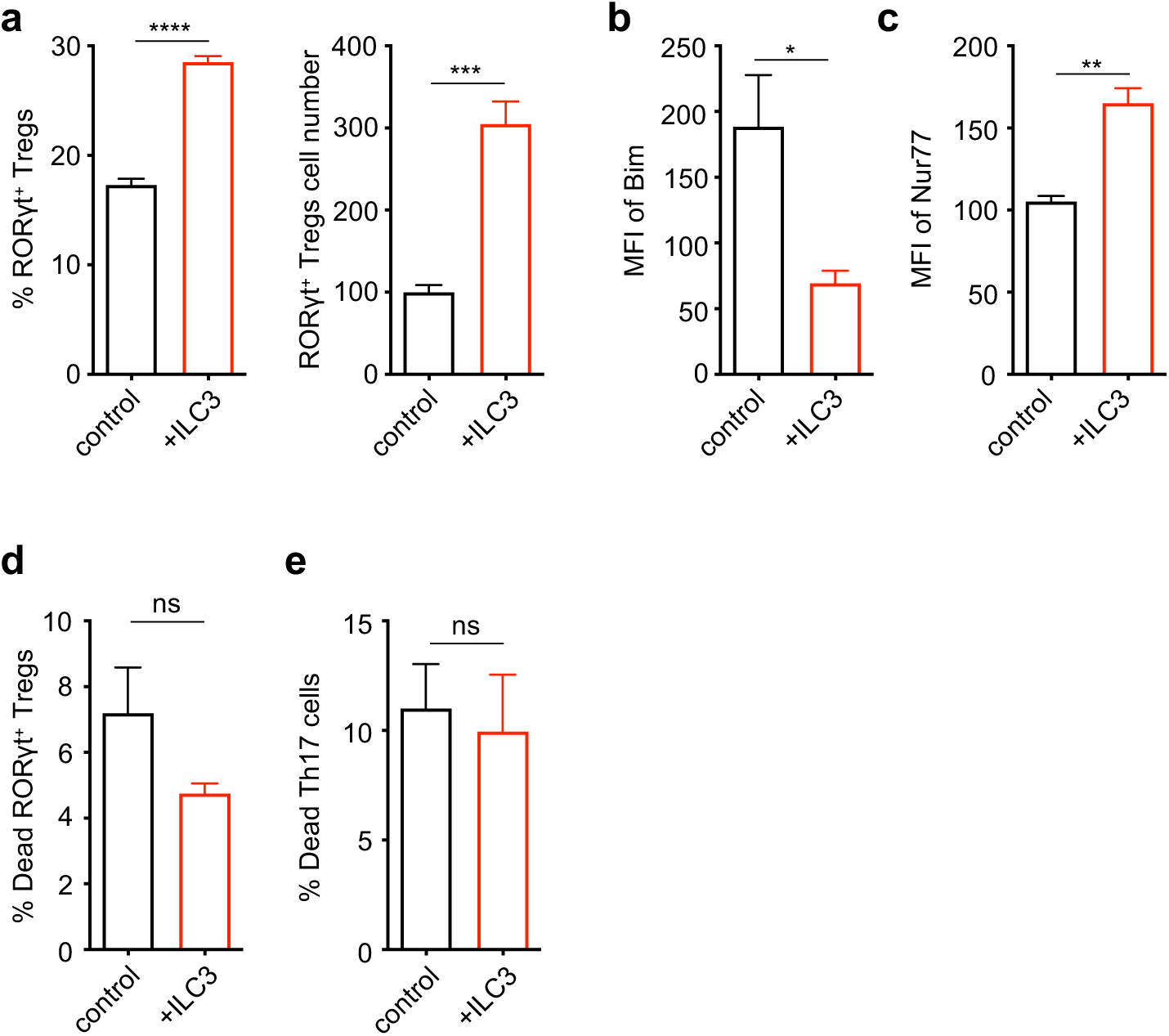
LTi-like ILC3s support RORγt^+^ Tregs in a co-culture system. **a**-**c**, RORγt^+^CD4^+^ T cells and LTi-like ILC3s were co-cultured for 72 hours and RORγt^+^ Tregs were analyzed by flow cytometry. Frequency and cell number of RORγt^+^ Tregs (**a**, RORγt^+^Foxp3^+^ among CD4^+^ T cells), MFI of Bim (**b**) and Nur77 (**c**) in RORγt^+^ Tregs. **d**, **e**, Dead cells were quantified in RORγt^+^ Tregs (**d**) and Th17 cells (**e**) after co-culture with or without LTi-like ILC3s for 72 hours. RORγt^+^CD4^+^ T cells and LTi-like ILC3s were sort-purified from mLN and LI-LP and pooled for co-culture assay. Data in (**a**-**c**) are representative of two independent experiments. Data in (**d**, **e**) are pooled from two independent experiments. Data are shown as mean ± SEM. Statistics were obtained by unpaired Student’s *t*-test (two-tailed). ns, not significant; *p<0.05, **p<0.01, ***p<0.001, ****p<0.0001.

**Extended Data Fig. 8.**
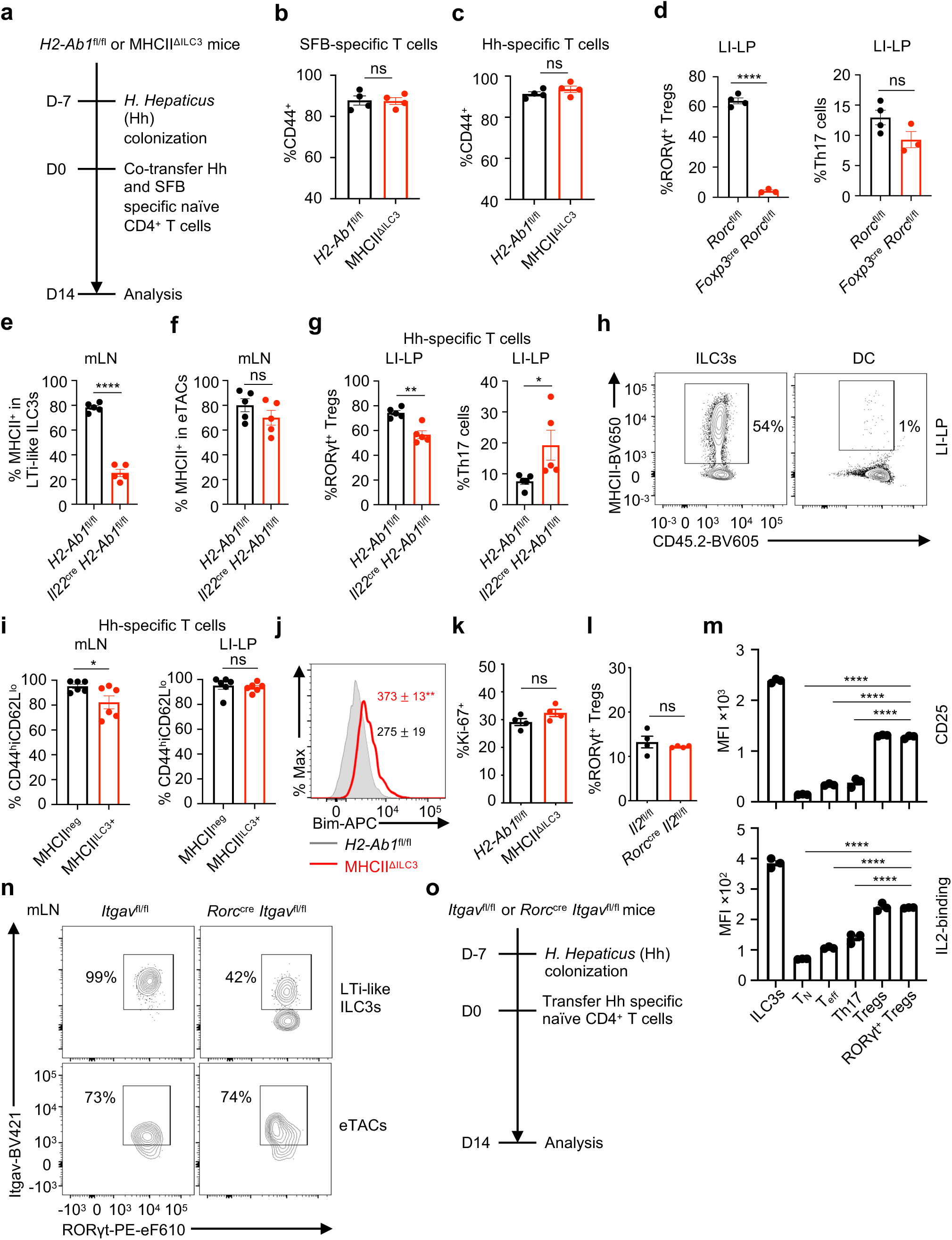
MHCII and Itgav on LTi-like ILC3s contributes to the selection of microbiota-specific RORγt^+^ Tregs. **a**, *H. Hepaticus* (Hh)-specific (Hh7-2) and/or SFB-specific (SFB7B8) CD4^+^ T cells were transferred to *H2-Ab1*^fl/fl^ and MHCII^ΔILC3^ mice colonized with *H. Hepaticus* 2 weeks before experiment as shown in Fig. 3 a-h, k, l, Fig. 4f. **b**, **c**, Frequency of CD44^+^ ratio among SFB-specific (**b**) or Hh (**c**) CD4^+^ T cells were analyzed in Peyer’s patch for SFB7B8 (CD45.1^-^CD90.1^+^CD4^+^ T cells) and in colon for Hh7-2 (CD45.1^+^CD90.1^-^CD4^+^ T cells) (n = 4). **d**, Quantification of RORγt^+^ Tregs and Th17 cells among total CD4^+^ T cells in LI-LP of *Rorc*^fl/fl^ and *Foxp3*^cre^ x *Rorc*^fl/fl^ mice (n = 4). **e**, **f**, Quantification of MHCII expression on LTi-like ILC3s (**e**) and eTACs (**f**) in mLN of *H2-Ab1^fl/fl^* and *Il22^cre^* x *H2-Ab1^fl/fl^* mice (n = 5). **g**, Quantification of RORγt^+^ Tregs (RORγt^+^FoxP3^+^ among Hh-specific CD4^+^ T cells, left) and Th17 cells (RORγt^+^FoxP3^-^ among Hh-specific CD4^+^ T cells, right) were analyzed in LI-LP of *H2-Ab1^fl/fl^* and *Il22^cre^* x *H2-Ab1^fl/fl^* mice (n = 5). **h**, Representative flow cytometry plots of the frequency of MHCII expression on ILC3s, DCs in LI-LP of MHCII^neg^ and MHCII^ILC3+^ mice (n = 6) as shown in Fig. 3j. **i**, Frequency of CD44^hi^CD62L^lo^ ratio among Hh CD4^+^ T cells were analyzed in mLN and LI-LP for Hh7-2 as shown in Fig. 3k, l (n = 4). **j**, **k**, RORγt^+^ Tregs of *H2-Ab1*^fl/fl^ and MHCII^ΔILC3^ mice large intestines were analyzed by flow cytometry. Histogram and MFI of Bim (**j**). Proportions of Ki-67 positive cells (**k**). **l**, Quantification of RORγt^+^ Tregs (RORγt^+^Foxp3^+^ among CD4^+^ T cells) in large intestine of *Il2*^fl/fl^ and *Rorc*^cre^*Il2*^fl/fl^ mice were analyzed (n = 4). **m**, Quantification of CD25 staining or IL-2 binding in mLN in WT mice (n = 3). Naive T cells were gated as CD44^lo^CD62L^hi^, effector T cells were gated as CD44^hi^CD62L^lo^, Th17 cells were gated as RORγt^+^FoxP3^-^, Tregs were gated as FoxP3^+^ and RORγt^+^Tregs were gated as RORγt^+^FoxP3^+^. **n**, Representative flow cytometry plot of the frequency of Itgav expression on LTi-like ILC3s and eTACs in mLN of *Itgav*^fl/fl^ and *Rorc*^cre^ x *Itgav*^fl/fl^ mice as shown in Fig. 4d (n = 5). **o**, *H. Hepaticus* (Hh)-specific (Hh7-2) CD4^+^ T cells were transferred to *Itgav*^fl/fl^ and *Rorc*^cre^ x *Itgav*^fl/fl^ mice colonized with *H. Hepaticus* 14 days before experiment related to Fig. 4f. Data are representative of two independent experiments. Data are shown as mean ± SEM, statistics shown in (**m**) are obtained by one-way ANOVA with Tukey’s multiple comparisons test, statistics shown in (**b**-**g**, **i**-**l**) are obtained by unpaired Student’s *t*-test (two-tailed), ns, not significant; *p<0.05, **p<0.01, ****p<0.0001.

**Extended Data Fig. 9.**
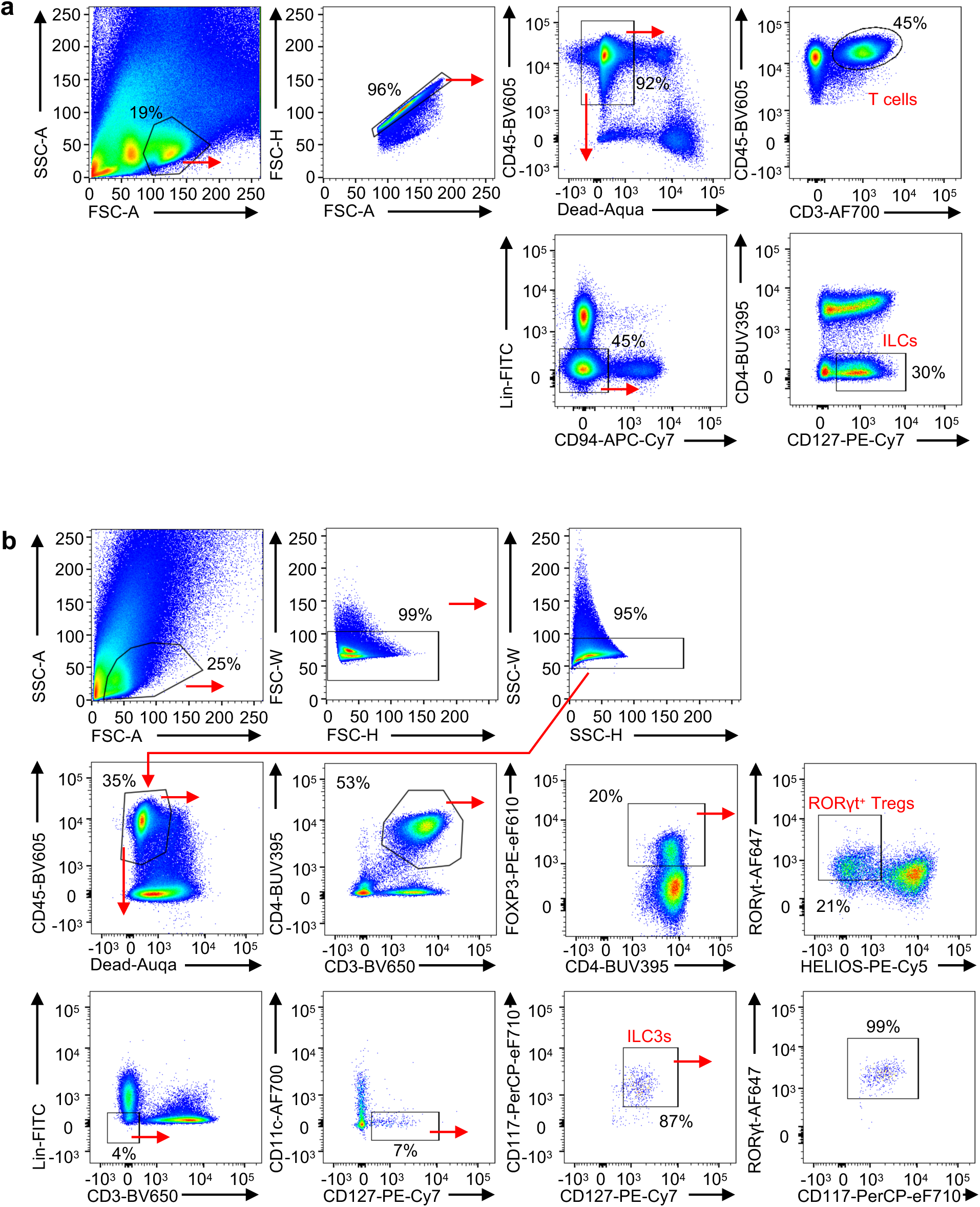
Gating strategy for ILCs and T cells in the human intestine. **a**, Gating strategy to sort ILCs and T cells from small intestine of the IBD patient for scRNA-seq. **b**, Gating strategy to demonstrate the ILC3s and RORγt^+^ Tregs shown in Fig. 5f, k, l.

**Extended Data Fig. 10.**
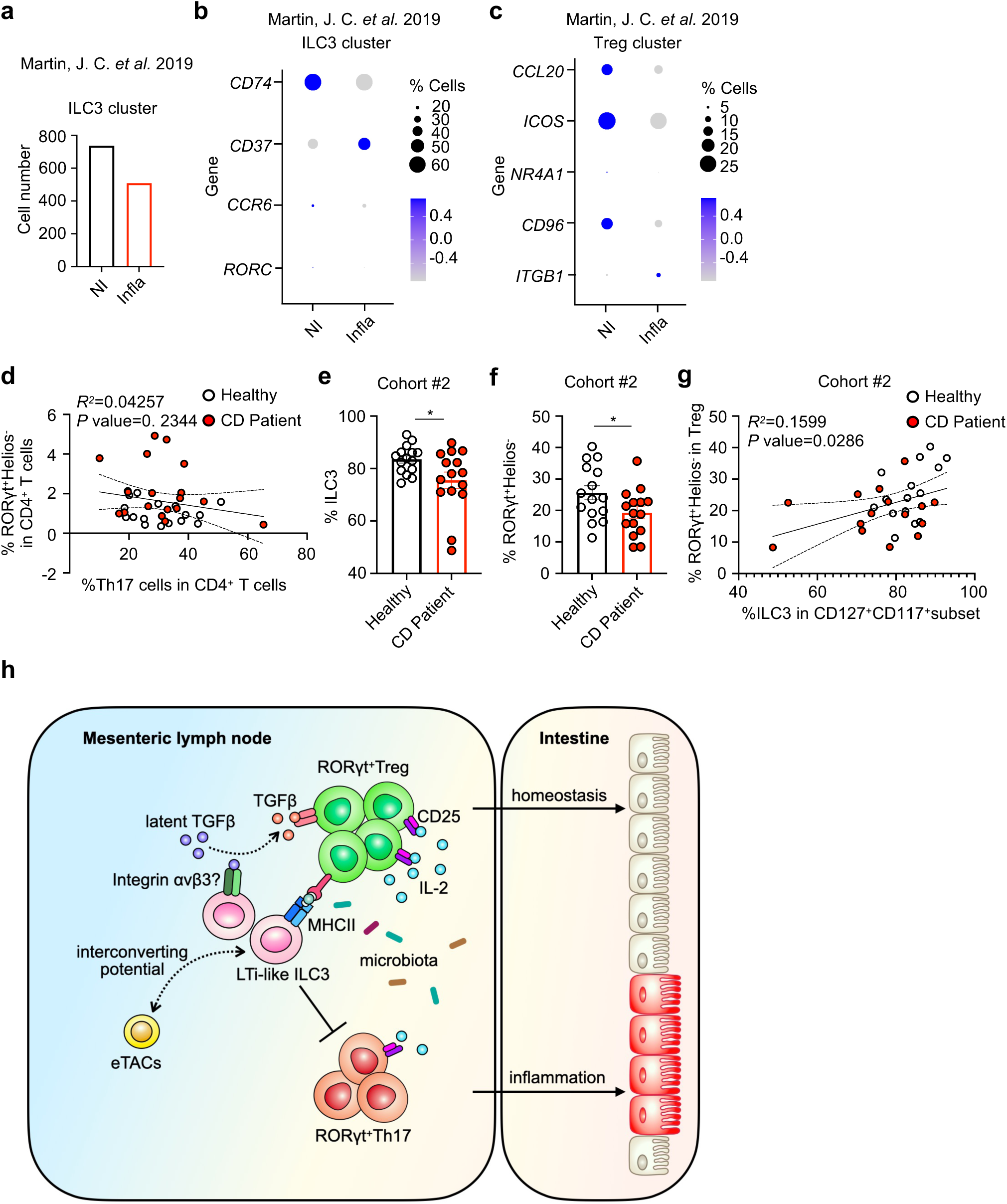
ILC3s select for RORγt^+^ regulatory T cells and enforce tolerance to intestinal microbiota. **a**, Bar graph showing the composition of ILC3 lymphocytes in non-inflamed tissue (NI) versus inflamed tissue (Infla) from patient as published^44^. **b**, **c**, Expression of indicated genes in ILC3 cluster (**b**) and Treg cluster (**c**) in non-inflamed versus inflamed tissue from patient as published^44^. **d**, Correlation analyses between the RORγt^+^ Tregs (RORγt^+^Helios^-^ among CD4^+^ T cells) and Th17 cells (RORγt^+^FoxP3^-^ among CD4^+^ T cells) in the cohort of CD patients as in Fig.5f, k, l. **e, f,** Quantification of frequency of ILC3 among CD127^+^CD117^+^ subset (**e**) and RORγt^+^ Tregs among total Tregs (**f**) in a second cohort of Crohn’s disease (CD) patients. Healthy donor n = 15, CD patient n = 15. **g**, Correlation analyses between the ILC3 (ILC3 among CD127^+^CD117^+^ subset) and RORγt^+^ Tregs (RORγt^+^ Helios^-^ among FoxP3^+^ Tregs) in a second cohort of CD patients as in (**e**, **f**). **h**, LTi-like ILC3s are indispensable for selecting for the differentiation fates of microbiota-specific RORγt^+^ Tregs, and against Th17 cells, via antigen presentation with contributions from integrin αv and gradients of competition for IL-2. This collectively enforces immunologic tolerance to microbiota and maintains intestinal homeostasis. Data in (**e**, **f**) are shown as means ± SEM, statistics shown in (**e**, **f**) are performed using Mann– Whitney U-test (unpaired), correlative analyses in (**d, g**) are compared by Pearson’s rank correlation coefficient (*R^2^*), *p<0.05.

